# PRC2 Inactivating Mutations in Cancer are Synthetic Lethal with DNMT1 Targeted Therapy via Enhanced Viral Mimicry

**DOI:** 10.1101/2022.05.28.493429

**Authors:** Amish J. Patel, Sarah Warda, Jesper L.V. Maag, Rohan Misra, Miguel A. Miranda-Román, Mohini R. Pachai, Dan Li, Naitao Wang, Cindy J. Lee, Gabriella Bayshtok, Eve Fishinevich, Yinuo Meng, Elissa W.P. Wong, Juan Yan, Emily Giff, Melissa B. Pappalardi, Michael T. McCabe, Jonathan A. Fletcher, Joseph M. Scandura, Richard P. Koche, Jacob L. Glass, Cristina R. Antonescu, Deyou Zheng, Yu Chen, Ping Chi

## Abstract

Polycomb Repressive Complex 2 (PRC2) establishes and maintains di- and tri-methylation at histone 3 at lysine 27 (H3K27me2/3) in the genome and plays oncogenic and tumor suppressor roles in context-dependent cancer pathogenesis. While there is clinical success of therapeutically targeting PRC2 core component, EZH2, in PRC2-dependent cancers (e.g., follicular lymphoma, epithelioid sarcoma), it remains an unmet therapeutic bottleneck in PRC2-inactivated cancer. Biallelic inactivating mutations in PRC2 core components are a hallmark feature of high-grade malignant peripheral nerve sheath tumor (MPNST), an aggressive subtype of sarcoma with poor prognosis and no effective targeted therapeutics. Using a custom RNAi-based drop out screen, we observed that PRC2-inactivation is synthetic lethal with DNA methyltransferase 1 (DNMT1) downregulation; we further observed that small molecule DNMT inhibitors (DNMTis) resulted in enhanced cytotoxicity and antitumor response in PRC2-loss cancer context *in vitro* and *in vivo*. Mechanistically, DNMTi-mediated de-repression of retrotransposons (e.g., endogenous retroviral elements (ERVs)/LTR, LINE, SINE) and gene targets is partly restricted by PRC2, which potentially contributes to limited therapeutic activity in PRC2-wild-type (wt) cancer context. In contrast, DNMTi treatment synergizes with PRC2 inactivation and cooperatively amplifies the expression of retrotransposons (e.g., ERV/LTR, LINE, SINE), and subsequent viral mimicry response that promotes robust cell death in part through PKR-dependent double stranded-RNA (dsRNA) sensing. Collectively, our observations posit DNA methylation as a safeguard against anti-tumorigenic cell fate decisions in the context of PRC2-inactivation to promote cancer pathogenesis. Further, they identified a novel targeted therapeutic strategy in PRC2-inactivated MPNST and delineated the PRC2-inactivated cancer context for future preclinical exploration and clinical investigation of DNMT1-targeted therapies in cancer.

**SIGNIFICANCE:** PRC2-inactivation drives oncogenesis in various cancers but therapeutically targeting PRC2-loss has remained challenging. Here we show that PRC2 inactivating mutations sets up a tumor context-specific liability for synthetic lethal interaction with genetic and therapeutic inhibition of DNMT1. DNMT1 inhibitor-induced cytotoxicity in PRC2-loss cancer context is accompanied by innate immune signaling signature through PKR-mediated sensing of endogenous retrotransposons. These observations posit a therapeutic window via direct anti-tumor effect by DNMT1 inhibitors in PRC2-loss cancers, and point to potentials to be combined with innovative immunotherapeutic strategies to capitalize on innate immune signaling activation.

## INTRODUCTION

The PRC2 complex consisting of core components SUZ12, EED, and EZH1/2, establishes and maintains H3K27me2/3 throughout the genome and regulates chromatin structure, transcription, cellular stemness and differentiation^1^. In cancer pathogenesis, PRC2 can function as an oncogenic driver through *EZH2* overexpression in various cancers^2,3^ and gain-of-function *EZH2* mutations in lymphomas^4^; PRC2 can also function as a tumor suppressor through inactivating genetic alterations in myeloid disorders^5,6^, T-cell acute lymphoblastic leukemia (ALL)^7^, early T-cell precursor ALL^8^, melanoma^9^, and malignant peripheral nerve sheath tumor (MPNST)^9-11^, or by functional inactivation via onco-histone mutations (H3K27M) in pediatric gliomas^12^. Complete inactivation of PRC2 is most prevalent, occurring in ∼80% of high-grade MPNSTs^13^.

MPNST represents an aggressive subtype of soft tissue sarcoma with poor prognosis due to the surgical challenges in local control and lack of effective systemic therapy. They occur in distinct clinical settings: type I Neurofibromatosis (NF1)-associated (45%), sporadic (45%) or radiation (RT)-associated (10%), yet share highly recurrent and biallelic genetic inactivation of three tumor suppressor pathways: *NF1, CDKN2A*, and PRC2 (*EED* or *SUZ12*)^14-16^. PRC2 inactivation results in global loss of H3K27me2/3 and aberrant transcriptional activation of developmentally silenced master regulators, leading to enhanced cellular plasticity in MPNST^15,17,18^. Therapeutically targeting PRC2-inactivation in cancer remains a challenge and requires relevant context specific cancer models for therapeutic discovery and development.

To discover synthetic lethal interactions with PRC2 inactivating alterations in their relevant cancer setting, we conducted an epigenome-focused pooled RNAi dropout screen in patient-derived MPNST cell lines. We identified and validated DNMT1, whose down-regulation resulted in selective growth inhibition in PRC2-loss MPNST *in vitro* and *in vivo*. Therapeutic targeting of DNMT1 with the FDA approved Decitabine (a pan-DNMT inhibitor) or a novel preclinical DNMT1-selective catalytic inhibitor led to significantly enhanced cytotoxicity dependent on the loss of PRC2 enzymatic activity. Mechanistically, PRC2 and DNA methylation coregulate transcriptional silencing of a subset of H3K27me3-enriched transcriptional targets and retrotransposons. Loss of PRC2 function amplifies DNMTi therapy induced transcriptional activation of retrotransposons and subsequent viral mimicry mediated cell death through dsRNA sensor PKR. These data point to PRC2 inactivating mutations as a driver of cancer pathogenesis, and a liability that primes tumor cell-selective and robust cell death via DNMT1 targeted therapy while sparing PRC2-wild type (wt) normal cells.

## RESULTS

### RNAi screening identifies *DNMT1* synthetic lethality with PRC2 inactivation in cancer

To identify epigenetic factors that pose selective synthetic lethal interactions with PRC2-inactivation, we conducted a pooled doxycycline-inducible short hairpin RNA (shRNA) screen targeting 565 known epigenetic/chromatin regulators in PRC2-wt and PRC2-loss patient-derived MPNST cell lines. We authenticated that all cell lines have complete loss of both *NF1* and *CDKN2A*, and 5 of 7 (∼71%) cell lines had complete PRC2 inactivation by MSK-IMPACT^19^ and/or immunoblotting (**Fig. 1A** and **Supplementary Fig. S1**). After shRNA screening (**Fig. 1B**), we used HiTSelect^20^ to rank each set of 5 shRNAs per gene in the library across all cell lines in the screen for depletion (negative selection) (**Supplementary Table 1**). As anticipated, shRNAs targeting an essential gene (e.g. *ACTB*) were ranked among the top (most significant) while non-toxic control shRNA (sh*Ren*.713) was ranked at the bottom (least significant) across all cell lines (**Fig. 1C**). Among all candidates, *DNMT1*-specific shRNAs were ranked consistently among the top 10 statistically significant candidates for negative selection across all PRC2-loss but not PRC2-wt cell lines (**Fig. 1C**). *DNMT1* encodes a DNA methyltransferase responsible for maintaining DNA methylation (5 - methylcytosine) at CpG dinucleotides on newly replicated DNA during cellular division^21^. Interestingly, shRNAs specific for *de novo* DNA methyltransferases (*DNMT3A* and *DNMT3B*), were less significantly ranked in the screen (**Fig. 1C**). We found that *DNMT1* is consistently and abundantly expressed in all human MPNST cell lines irrespective of their PRC2 status, while *DNMT3A* and *DNMT3B* expression is variable without any correlation to PRC2 status (**Supplementary Fig. S2A-B**). These data suggest that the selective DNMT1-PRC2 loss synthetic lethal interaction is through mechanisms distinct from selective *DNMT1* expression in MPNST tumor cells.

**Figure 1.**
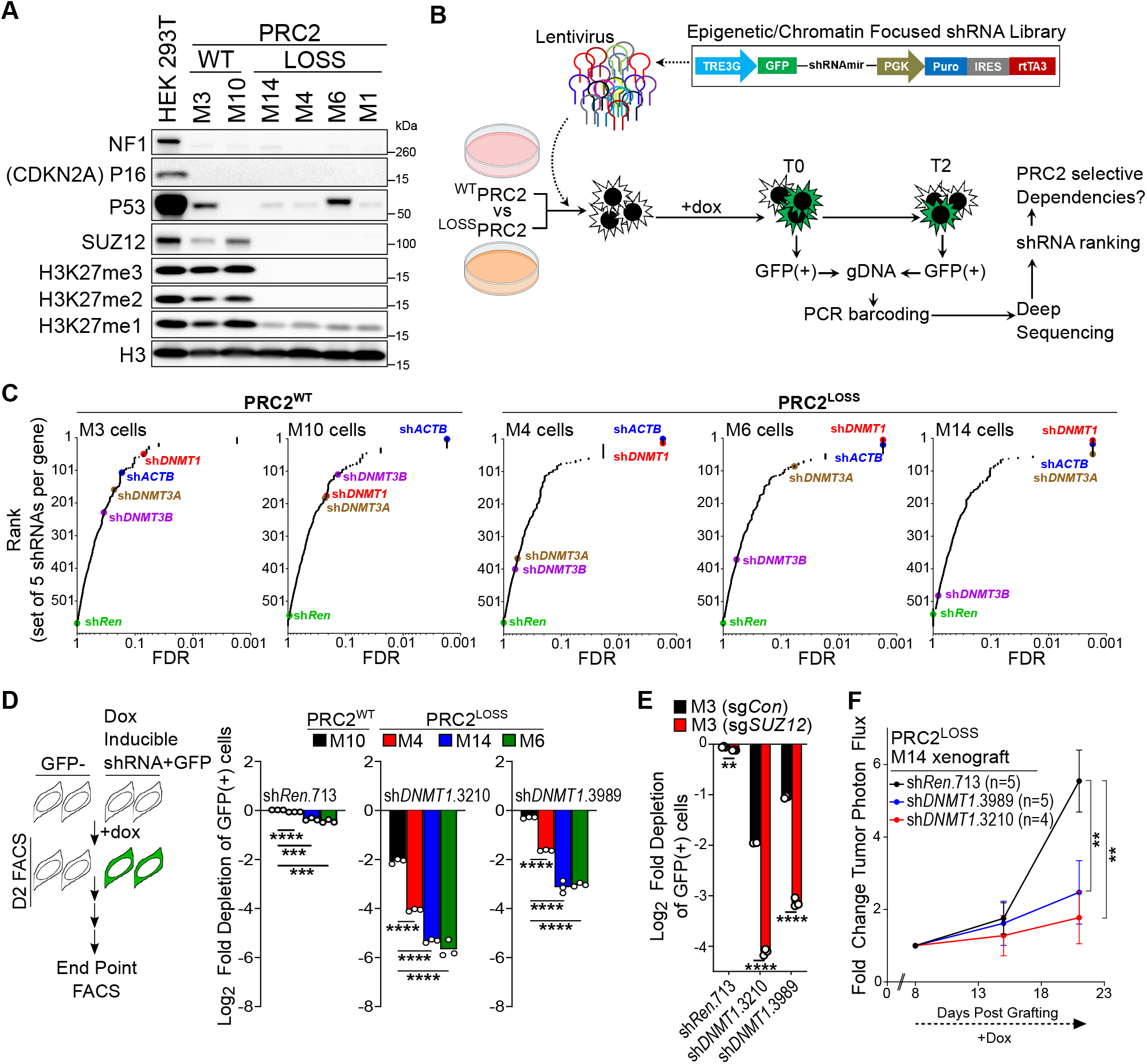
RNAi screen identifies DNMT1 as a synthetic lethal target in PRC2-inactivated MPNST. **A**, Western blot validation of NF1, CDKN2A, and PRC2 status in human MPNST cell lines. **B**, Schematics of the RNAi-based pooled dropout screen in PRC2-wt (M3, M10) and PRC2-loss (M4, M6, M14) MPNST cell lines. **C**, shRNA gene-set rank and statistical significance (FDR) plot based on HiTSelect algorithm for shRNA depletion in each MPNST cell line. **D**, Growth competition assay by FACS using shRNA-linked GFP under DNMT1-specific perturbations (sh*DNMT1*.3210, sh*DNMT1*.3989) compared to non-targeting control (sh*Re*n.713) in MPNST cell lines. **E**, Validation of *DNMT1* as a synthetic lethal candidate with PRC2-inactivation using shRNA-linked GFP FACS-based growth competition assay in PRC2-isogenic MPNST cells (PRC2-wt = M3[sg*Con*] vs. PRC2-loss = M3[sg*SUZ12*]). **F**, Luciferase bioluminescence-based growth curve over time for orthotopic human MPNST xenografts with doxycycline (dox) inducible knockdown of *DNMT1*. All error bars: mean ± SEM; Student’s two-tailed T-test (\*\**P*<0.01; \*\*\**P*<0.001; \*\*\*\**P*<0.0001).

To validate whether PRC2 is a mediator of DNMT1 dependency in MPNST, we used two different *DNMT1*-specific shRNAs from the screen that efficiently depleted *DNMT1* mRNA expression across multiple MPNST cell lines (**Supplementary Fig. S2C**). We monitored the effect of shRNAs by a flow cytometry (FACS)-based growth competition assay between GFP positive-shRNA expressing versus GFP negative-shRNA non-expressing MPNST cells, and observed significantly more rapid depletion of GFP positive-*DNMT1* shRNA expressing cells in PRC2-loss compared to PRC2-wt MPNST cell lines (**Fig. 1D**). To confirm that enhanced sensitivity to *DNMT1* depletion is dependent on PRC2 status, we generated PRC2-isogenic human MPNST cells via CRISPR/Cas9-mediated knockout of *SUZ12* (sg*Con* [PRC2-wt] vs. sg*SUZ12* [PRC2-loss]) (**Supplementary Fig. S2D-E**). Knockout of *SUZ12* did not markedly affect the protein expression of DNMT1, DNMT3A, or DNMT3B (**Supplementary Fig. S2D-E**). Importantly, we observed enhanced sensitivity to *DNMT1*-specific, but not *DNMT3A/B*-specific knockdown in PRC2-loss compared to PRC2-wt MPNST isogenic background (**Fig. 1E** and **Supplementary Fig. S2F-J**). To evaluate the cross-dependency *in vivo*, we orthotopically transplanted luciferase-tagged PRC2-loss human MPNST cells stably expressing doxycycline-inducible shRNAs specific for *DNMT1*. We observed that *DNMT1* shRNAs markedly suppressed MPNST tumor growth (**Fig. 1F**). These data confirmed the selective vulnerability of PRC2-inactivation to *DNMT1* depletion in cancer cells and tumors.

### DNMT1 targeted therapy exhibits selective anti-tumor activity against PRC2-loss cancer

The RNAi screening results compelled us to evaluate whether the selective dependence on DNMT1 in PRC2-loss MPNST can be leveraged therapeutically. DNMT1 can be inhibited by a clinically available pan-DNMTi decitabine (DAC), which is a nucleoside analog that inhibits DNMT1, DNMT3A and DNMT3B upon incorporation into DNA during S-phase^22^. Alternatively, DNMT1 can be selectively and directly inhibited by a novel preclinical non-covalent small molecule enzymatic inhibitor GSK3484862 (referred to as GSK862 hereafter)^23^. By dose response analysis, we observed that PRC2 inactivation by CRISPR/Cas9-mediated knockout of *SUZ12* enhanced sensitivity to both DNMTis (**Fig. 2A-B** and **Supplementary Fig. S3A**). Long-term growth assays confirmed that DAC and GSK862 similarly exhibited enhanced cytotoxicity selective for PRC2-loss MPNST cells (**Supplementary Fig. S3B-C**). To further evaluate the role of cellular PRC2 activity on sensitivity to DNMTi therapy, we treated PRC2-isogenic MPNST cells with an EZH2 catalytic inhibitor (EPZ-6438)^24^ or an H3K27me3 binding pocket inhibitor of EED (EED226)^24,25^. EZH2 and EED inhibitors effectively reduced H3K27me2/3 in PRC2-isogenic M3 cells and PRC2-wt M10 cells (**Fig. 2C** and **Supplementary Fig. S3D**), and led to enhanced cytotoxicity upon treatment by DNMT1 inhibitor (GSK862) in PRC2-wt MPNST cells (M3-sgCon and M10), but not in PRC2-loss cells (M3-sg*SUZ12*) (**Fig. 2D** and **Supplementary Fig. S3E**).We further tested DNMTi therapy in established PRC2-isogenic (*Eed*^*-/-*^vs. *Eed*^f/-^) murine MPNST cells derived from genetically engineered mouse models of MPNST that also harbor complete loss of *Nf1* and *Cdkn2a* (**Fig. 2E** and **unpublished**). Similarly, we observed enhanced sensitivity to DNMTi in PRC2-loss (*Eed*^*-/-*^) compared to PRC2-intact (*Eed*^f/-^) murine MPNST cells (**Fig. 2E**). To further validate the selectivity of the DNMTi in PRC2-loss cells, we restored PRC2 activity by ectopically expressing *SUZ12* cDNA into *SUZ12*-null patient-derived MPNST cell lines, and confirmed PRC2 activity (H3K27me3 and repression of PRC2 target genes) (**Supplementary Fig. S3F-G**). We observed decreased sensitivity to DNMTi upon PRC2 restoration (**Fig. 2F**and **Supplementary Fig. S3H)**, thus corroborating the PRC2 and DNMT pathway cross-dependence in MPNST. To evaluate the effect of DNMTi therapy *in vivo*, we treated various patient-derived MPNST xenograft tumor models with the pan-DNMTi (DAC). We observed that DAC exhibits selective anti-tumor activity against PRC2-loss tumors (**Fig. 2G-I**). To specifically inhibit DNMT1 *in vivo*, we utilized GSK3685032, a DNMT1-selective inhibitor that is structurally related to GSK862, but capable of *in vivo* inhibition of DNMT1 in mice^23^. Consistent with GSK862 (**Fig. 2F**), GSK3685032 effectively inhibits *in vitro* growth of PRC2-loss MPNST cells, but not when *SUZ12* is re-expressed (**Supplementary Fig. S3I)**.

**Figure 2.**
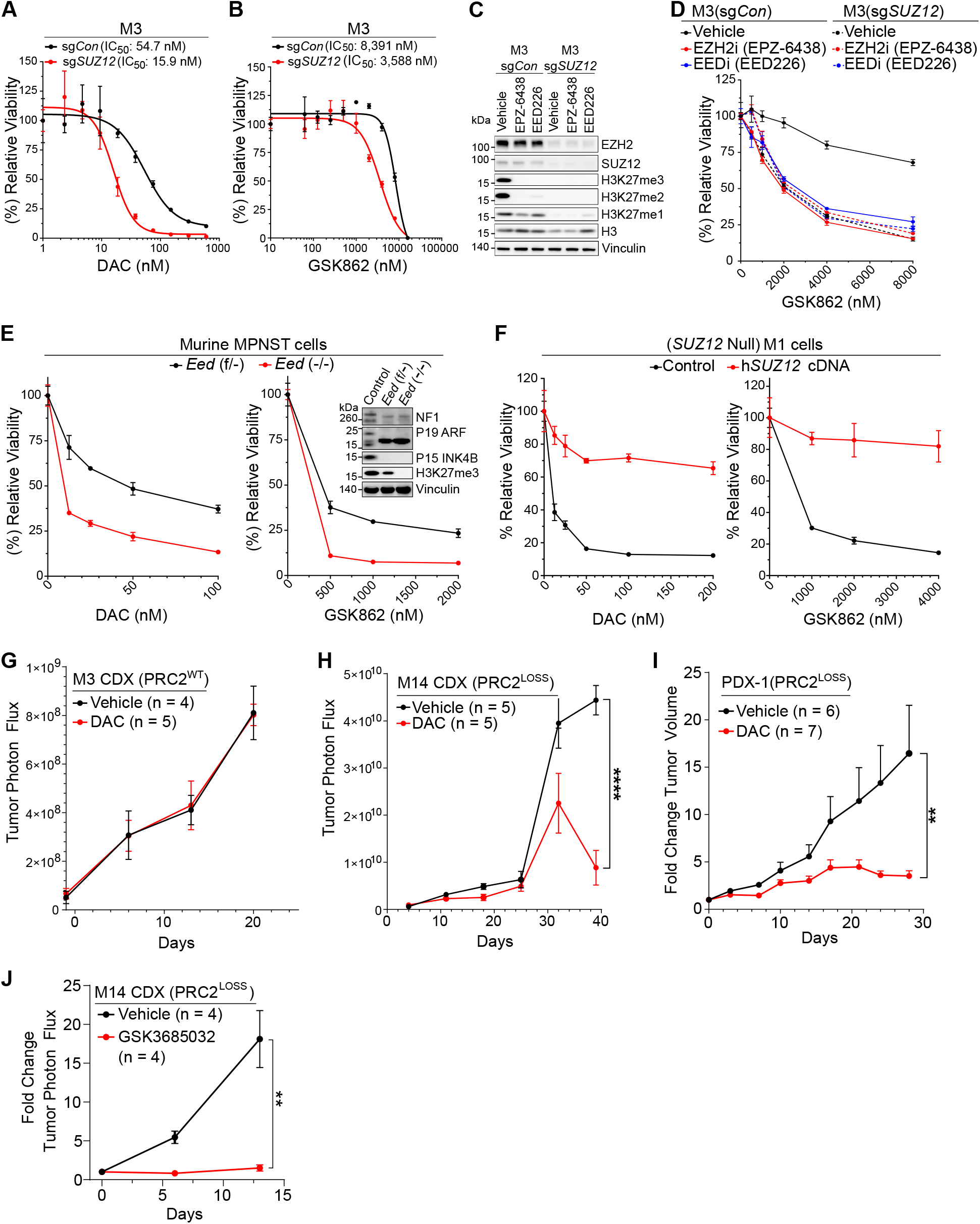
DNMT1 targeted therapy has selective anti-tumor activity for PRC2-loss cancer. **A-B**, Dose-response curves and IC_50_ of a pan-DNMT inhibitor, decitabine (DAC), or a DNMT1-selective inhibitor (GSK862) in *SUZ12*-isogenic MPNST cells: PRC2-wt= M3-sg*Con*; PRC2-loss = M3-sg*SUZ12*. **C**, Western blot analysis of *SUZ12-*isogenic cells treated with 3 μM EPZ-6438 (an EZH2 inhibitor) or 5 μM EED226 (an EED inhibitor). **D**, Dose-response curves of GSK862 in *SUZ12-*isogenic cells treated with EZH2 or EED inhibitors, or vehicle controls. **E**, Western blot validation and DNMT inhibitor (DAC and GSK862) dose-response curves for *Eed-*isogenic murine MPNST cells derived from *Nf1*^-/-^;*Cdkn2a*^-/-^;*Cdkn2b*^*-/-*^ genetically engineered murine models. **F**, Dose-response curves of DAC and GSK862 in *SUZ12*-null human MPNSTs cells with or without PRC2 (*SUZ12*) restoration. **G-H**, Growth curves over time of orthotopically transplanted human PRC2-wt (M3) and PRC2-loss (M14) cell-line-derived xenografts (CDXs) in CB-17 SCID mice treated with DAC or vehicle control by luciferase bioluminescence imaging. **I**, Fold change tumor volume of subcutaneous PRC2-loss human MPNST patient derived xenograft (PDX-1) treated with 5 mg/kg DAC or vehicle control. **J**, Luciferase bioluminescence-based growth curve over time of orthotopically transplanted human PRC2-loss M14 CDX in SCID mice treated with 45 mg/kg GSK3685032 or vehicle. All error bars: mean ± SEM; Student’s two-tailed T-test (\*\**P*<0.01; \*\*\**P*<0.001; \*\*\*\**P*<0.0001).

Importantly, we observed that GSK3685032 potently inhibited orthotopic tumor growth of a PRC2-loss human MPNST CDX (**Fig. 2J**). Collectively, these observations indicate that PRC2 inactivation selectively sensitizes MPNST cells and tumors to DNMT pathway inhibition.

### CGI DNA methylation restricts transcription of a subset of PRC2/H3K27me3 target genes

Next, we investigated the molecular mechanisms underlying enhanced sensitivity to DNMT1 targeted therapy in PRC2-loss MPNST. We employed RNA-seq, bisulfite-seq, and ChIP-seq to characterize and integrate transcriptome, methylome, and histone modification responses to DNMT1 targeted therapy in PRC2-isogenic MPNST cells. *K*-means clustering analysis of differentially expressed genes under different conditions in RNA-seq transcriptomes revealed 6 distinct clusters of genes (**Fig. 3A**). Reasoning that PRC2 and DNMT1 are traditionally known for their roles in transcriptional silencing, we focused on genes up-regulated in clusters C1, C4, and C6 (**Fig. 3A**). As expected, C1 genes exclusively up-regulated by PRC2 loss were enriched for PRC2-related gene sets; C4 genes up-regulated solely by DNMTi treatment led to enrichment of previously published DNMTi-responsive gene sets (**Fig. 3B**). C6 genes were highly enriched for gene sets containing genes that have high density CpG promoters (HCP) and concurrent promoter H3K27me3 (**Fig. 3B**), which may suggest that C6 genes are co-regulated by PRC2 and DNA methylation. In contrast, genes with intermediate- and low-density CpG promoters (ICP and LCP) and concurrent H3K27me3 were marginally enriched in C6 (**Fig. 3B**). ChIP-seq analysis of PRC2-wt MPNST cells confirmed that a large subset of C1 and C6 genes have promoter H3K27me3 enrichment at baseline that is unaffected by DNMTi treatment (**Fig. 3C-D**). At the transcriptome level, most of the C1, C4 and C6 genes had low baseline mRNA expression regardless of H3K27me3 promoter enrichment, which is consistent with PRC2 and DNA methylation-mediated transcriptional silencing (**Fig. 3E**). Loss of PRC2 led to more robust transcriptional activation of C1 versus C6 genes (**Fig. 3E**). Similarly, DNMTi treatment led to more robust transcriptional activation of C4 versus C6 genes (**Fig. 3E**). The combined inactivation of PRC2 and DNMT pathways led to more robust transcriptional activation of C6 genes than by either perturbation alone (**Fig. 3E**).

**Figure 3.**
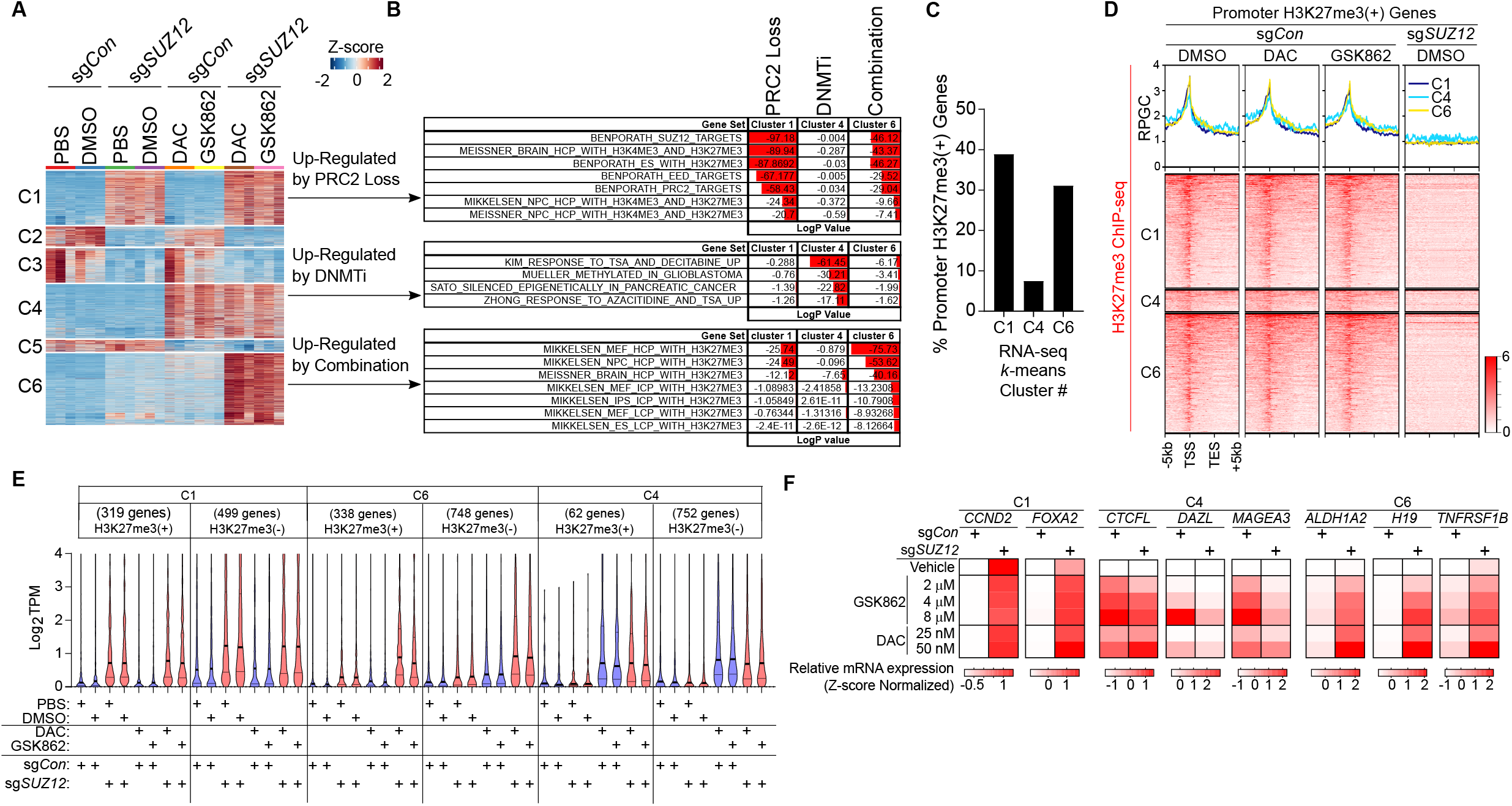
DNA methylation restricts transcription of a subset of PRC2-regulated genes. **A**, *k*-means clustering of differentially expressed genes for *SUZ12*-isogenic M3 MPNST cells. **B**, Gene set enrichment analysis identifies gene sets enriched by genes in RNA-seq *k*-means clusters 1 (C1), 4 (C4), and 6 (C6). **C**, Percentage of genes in C1, C4, and C6 that have a promoter enrichment of H3K27me3 by ChIP-seq in M3-sg*Con* cells. **D**, Enrichment profile of the average reads per genomic context (RPGC) (top) and heatmaps (bottom) of H3K27me3 ChIP-seq signals 5kb upstream of transcription start site (TSS) and 5kb downstream of transcription end site (TES) of H3K27me3(+) genes in C1, C4, C6 under various treatment conditions. **E**, Violin plot of RNA expression (Log2 TPM) of genes in C1, C4, and C6 separated by promoter H3K27me3 enrichment status (±) under various treatment conditions. Each violin has 3 lines inside. The middle thick line represents the median, and the upper and lower thin lines are the quartiles. **F**, Heatmap of RNA expression by qRT-PCR of representative genes in C1, C4, and C6 in *SUZ12*-isogenic M3 cells under indicated treatment conditions. Note: *CCND2, FOXA2, ALDH1A2, H19*, and *TNFRSF1B* each have promoter H3K27me3 enrichment in M3(sg*Con*) cells.

We confirmed transcript expression dynamics elicited by PRC2 and DNMT pathway inhibition under various conditions and across multiple MPNST model systems for representative genes in C1, C4, and C6. First, we confirmed that these trends occur in a dose-dependent manner in multiple PRC2-isogenic (sg*Con* vs. sg*SUZ12*) human MPNST cells *in vitro* (**Fig. 3F** and **Supplementary Fig. S4A**). Conversely, restoration of PRC2 activity dampened transcriptional activation of C6 genes by DNMTi therapy (**Supplementary Fig. S4B**). We confirmed that up-regulation of C1 and C6 gene expression is dependent on loss PRC2 enzymatic activity (**Supplementary Fig. S4C**). Moreover, these transcriptional changes were confirmed in PRC2-loss MPNST xenograft tumors *in vivo* with shRNA-mediated knockdown of *DNMT1* (**Supplementary Fig. S4D**). These data suggest that DNA methylation restricts a subset of PRC2-targets in C6.

We next investigated DNA methylation dynamics via capture-based enrichment of approximately 3 million representative CpGs coupled with bisulfite-sequencing (Methyl-seq) after PRC2 and DNMT pathway perturbation. We obtained DNA methylation data for 2,357,491 CpGs and summarized the CpG methylation data binned into 370,075 tiling windows (regions) across the genome (**Supplementary Fig. S5A**). Principal component analysis (PCA) demonstrated that both DNMTi therapies (GSK862 and DAC) caused significant DNA methylation changes regardless of PRC2 status; and DNMTi-treated samples were readily separated from vehicle-treated samples by principal component 1 (PC#1) (**Supplementary Fig. S5B**). The DNMTi-mediated DNA methylation changes were genome wide across all regions, including promoters, exons, introns, and CpG islands (CGIs) and shores (**Supplementary Fig. S5C**). In contrast, loss of PRC2 did not cause large-scale global changes in DNA methylation at regions we examined (**Supplementary Fig. S5C**). However, we identified a small number of differentially methylated regions (DMRs) (methylation difference > 25%, *q*< 0.01) as a result of PRC2 loss, either hypo- (n = 5,385) or hyper-methylated (n = 3,496) regions (**Supplementary Fig. S5D**). The percentage of hyper- and hypo-methylated DMRs were similar at each genomic and CpG regions (**Supplementary Fig. S5E-F**), and were associated with very few genes in RNA-seq *k*-means clusters C1, C4, and C6 (**Supplementary Data. S5G**). These data suggest that PRC2-loss mediated changes in DNA methylation is not the main mechanism underlying transcriptional silencing of C6 genes.

It is well established that DNA methylation at gene promoters is linked to repression of transcription initiation through various mechanisms (e.g. blockade of transcription factor binding, recruitment of corepressor complexes)^26^. We examined DNA methylation profiles spanning the transcription start site (TSS) to transcription end site (TES) of genes within each RNA-seq *k*-means cluster. As expected, the gene body and TES for all genes were DNA methylated while most of the active genes in clusters 2, 3, and 5 are DNA hypomethylated at their TSS/promoter regions (**Supplementary Fig. S5H**). Interestingly, genes preferentially upregulated by PRC2 loss (C1) had low levels of DNA methylation at their TSS/promoter regions; in contrast, genes preferentially upregulated by DNMTi (C4) or combined PRC2-loss and DNMTi (C6) had high or moderate levels of DNA methylation at their TSS/promoter regions, respectively (**Supplementary Fig. S5H**). We found a similar trend in DNA methylation profiles after separating genes in C1, C4, and C6 by H3K27me3 promoter enrichment status (**Fig. 4A**). Within the vicinity of the TSS, we consistently found that most C6 genes have elevated DNA methylation at their promoter, CGI, and shores compared to C1 genes (**Fig. 4B**).

**Figure 4.**
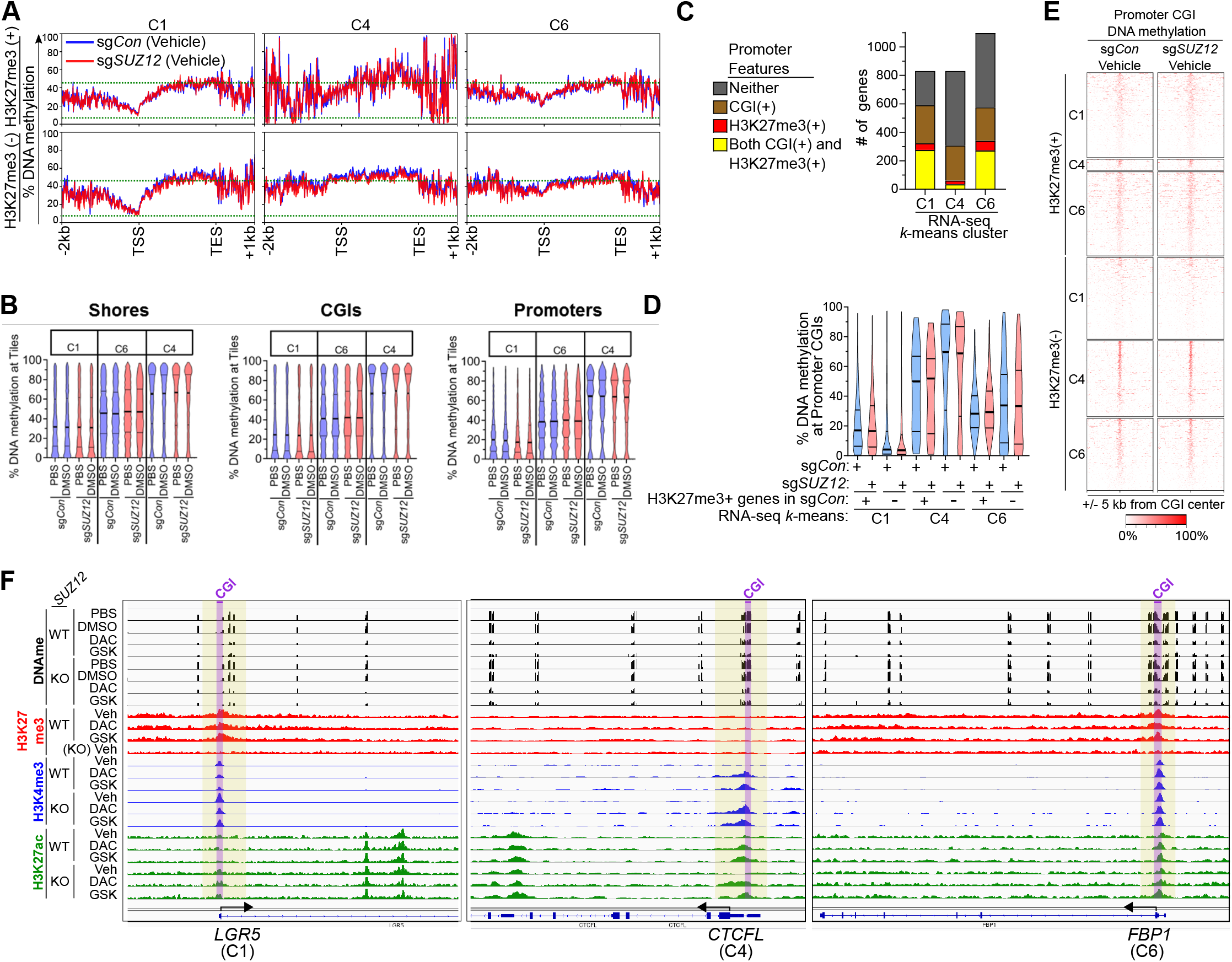
Promoter CGI DNA methylation associated with silencing of a subset of H3K27me3(+) and H3K27me3(-) genes in the absence of PRC2. **A**, Average DNA methylation profile plots spanning the TSS to TES for H3K27me3(+) and H3K27me3(-) genes in C1, C4, and C6 for M3(sg*Con*) and M3(sg*SUZ12*) cell lines. **B**, Violin plots of tile(region) DNA methylation levels for CpG shores, CGIs, and promoters of genes in C1, C4, and C6. **C**, Number of genes in specified RNA-seq *k*-means clusters that have a CGI and/or H3K27me3 enrichment at their respective promoters. **D**, Violin plots of DNA methylation levels within the promoter CGI coordinates based on H3K27me3 enrichment status(±) in specified RNA-seq *k*-means clusters in *SUZ12*-isogenic M3 cells. **E**, DNA methylation density heatmap for promoter CGI containing genes separated by H3K27me3 enrichment status(±)in C1, C4, and C6. **F**, ChIP-seq (H3K27me3, H3K4me3 and H3K27ac) and DNA methylation profiles of representative genes in C1, C4, and C6 in M3 *SUZ12*-isogenic cells (WT = sg*Con* ; KO = sg*SUZ12*). Promoter CGIs regions are shaded in purple.

Nearly ∼70% of known gene promoters contain CGIs that are CpG rich sites of transcription initiation when unmethylated, but transcriptionally repressed when densely methylated^26^. Moreover, H3K27me3 tends to be enriched at CGI sites (during *de novo* H3K27me3 methylation?)^27^. We found that ∼85% of H3K27me3(+) genes in C1 and ∼80% of H3K27me3(+) genes in C6 have a promoter CGI (**Fig. 4C**). We quantitatively and visually confirmed that most of the C6 promoter CGI containing genes had higher levels of DNA methylation compared to C1 genes (**Fig. 4D-E**). Inspection of genome tracks for representative H3K27me3(+) genes in C1 and C6 revealed that their promoter associated CGIs both have H3K27me3/H3K4me3 enrichment while DNA methylation is enriched for C6, but not C1 genes (**Fig. 4F**). This is consistent with dual modes of transcriptional repression observed for C6 genes at their promoters by PRC2 and DNA methylation. These data suggest that in the absence of PRC2, robust transcription of a subset of all H3K27me3 decommissioned target genes is restricted by the presence of CGI DNA methylation at their promoters.

### PRC2 inactivation amplifies DNMTi therapy induced viral mimicry

We next investigated the biological signature output for C6 genes, and its relevance to enhanced cytotoxicity to DNMTi therapy in PRC2-loss cancer context. We conducted gene ontology (GO) enrichment analysis for the six gene clusters in RNA-seq analysis and then performed hierarchical clustering of the GO terms to identify gene set signatures enriched by PRC2 loss and DNMTi co-regulated genes in cluster C6 (**Fig. 5A**). Immune gene sets related to innate immune response, interferon response, cytokines, and JAK/STAT signaling, as well as cell death signature gene sets were the most prominent (**Fig. 5A-B**). We confirmed that DNMTi therapy induced immune response signaling pathways (e.g., phosphorylation of TBK1 and STAT1) and expression of proinflammatory cytokines in MPNST cells, which is further enhanced by PRC2 loss (**Fig. 5C-D**). These were of interest because previous studies in other cancer types delineated that pan-DNMTi therapy triggers viral mimicry through de-repression of ERVs that form dsRNAs, which in turn primarily activate dsRNA sensor MDA5 to promote proinflammatory and antiviral innate immune response gene signatures ^28,29^. Aberrant expression of retrotransposons, including ERVs (LTR retrotransposons), SINEs (Short Interspersed Nuclear Elements), and LINEs (Long Interspersed Nuclear Elements) can all activate cytosolic dsRNA sensors and proinflammatory and innate immune responses in the absence of viral pathogen infection^30-35^. We therefore sought to determine if the ERV expression is regulated by PRC2 and DNA methylation. We specifically analyzed differentially expressed retrotransposons (e.g., ERVs, LINE, SINE) using RNA-seq derived from different PRC2 and DNA methylation perturbations (**Fig. 5E**). While there were subsets of retrotransposons upregulated by single perturbation of PRC2 loss or DNMTi, the ERVs were most pronouncedly and cooperatively upregulated by the combination of PRC2 loss and DNMTi treatment. These were confirmed by qRT-PCR in independent samples (**Fig. 5G**). These data suggest that PRC2 inactivation synergizes with DNMTi therapy in cancer cells and increase the expression of ERVs, SINEs, and LINEs to elicit a stronger viral mimicry transcriptional cell state than by DNMTi therapy or by PRC2 loss alone, which may contribute to the enhanced cytotoxicity of the combinatorial perturbations.

**Figure 5.**
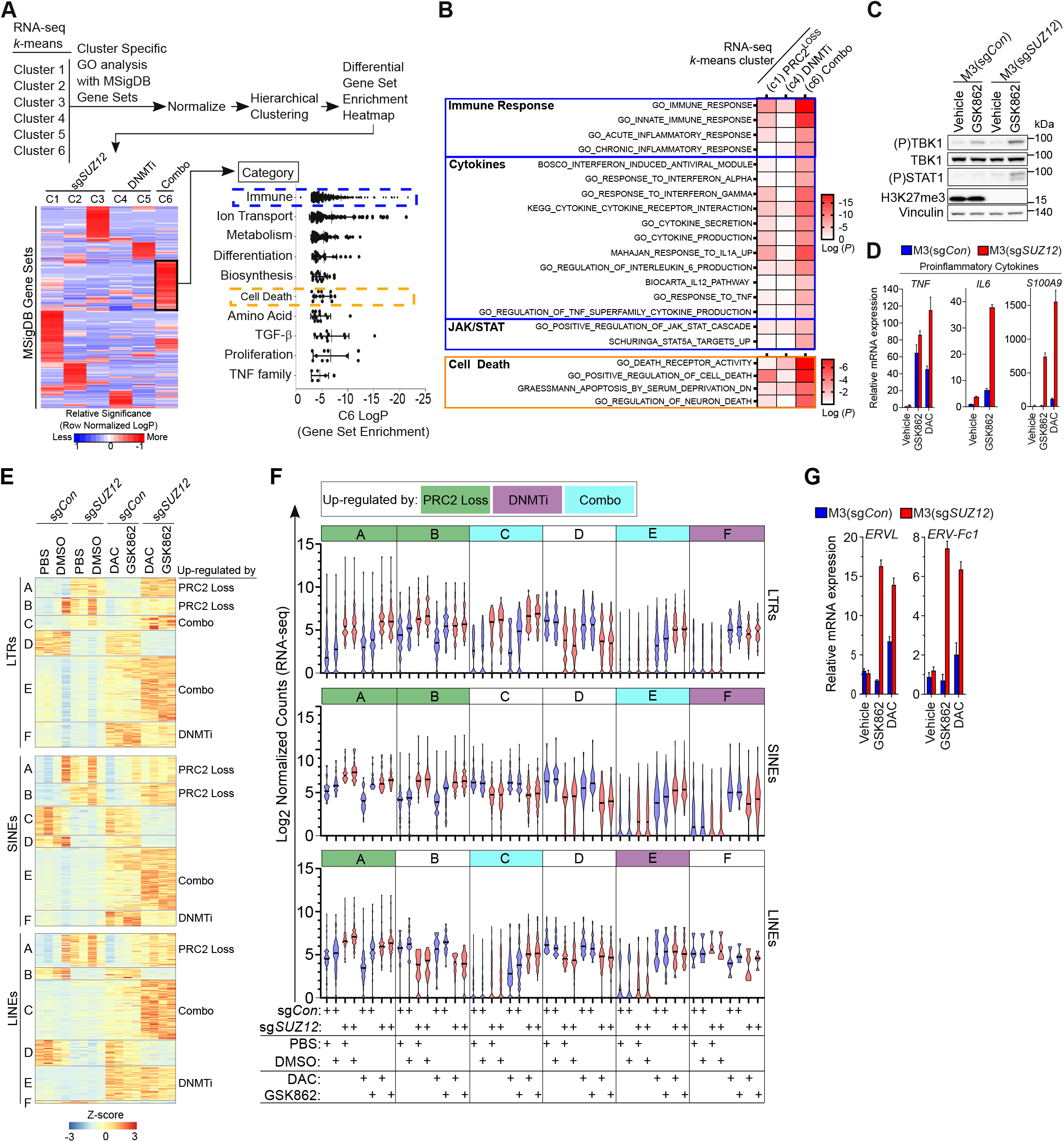
PRC2 inactivation potentiates DNMTi therapy induced viral mimicry and signaling response. **A**, Strategy for differential analysis of gene ontology (GO) terms enriched from the molecular signatures database (MSigDb) across each RNA-seq *k*-means cluster (from Figure 3a). Relative enrichment of MSigDb gene sets in each *k*-means cluster visualized by hierarchical clustering heatmap. Gene sets more enriched in cluster 6 were binned into different categories and plotted. **B**, Representative gene sets enriched preferentially by the combination of PRC2 loss and DNMTi therapy compared to single perturbation in *SUZ12*-isogenic M3 cells. **C**, Western blot analysis of viral mimicry signaling response (phosphorylation of TBK1 and STAT1) in *SUZ12*-isogenic cells treated with DNMTi therapy. **D**, qRT-PCR analysis proinflammatory cytokines in cells treated with DNMTi therapy. **E**, *k*-means clustering analysis for differentially expressed repetitive elements, including LTRs, SINEs, and LINEs in *SUZ12*-isogenic M3 cells treated under various conditions. **F**, Violin plot of log2 normalized RNA-seq counts for LTRS, SINEs, and LINEs differentially expressed in *SUZ12*-isogenic M3 cells under various treatment conditions. Each violin has 3 lines inside. The middle thick line represents the median, and the upper and lower thin lines are the quartiles. **G**, qRT-PCR analysis of ERV transcripts in cells treated with DNMTi therapy. All error bars: mean ± SEM.

### PRC2-loss selective cell death via DNMTi therapy is mediated by dsRNA sensor PKR

Historically, DNMTi (e.g., azacytidine or decitabine) is mostly cytostatic at clinically achievable concentrations and has limited activity in solid tumors^36,37^. Previous preclinical studies with pan-DNMTis where ERVs were derepressed also exhibited modest anti-proliferative activity^28,29^. In contrast, we noted increased enrichment of cell death gene signatures in response to DNMTi in PRC2-loss cancer context (**Fig. 5A-B**). We consistently observed that PRC2 inactivation in various cancer models converted a mostly cytostatic response into robust cytotoxicity upon DNMTi therapy via caspase-dependent programmed cell death (**Fig. 6A-D** and **Supplementary Fig. S6A-B**). These observations led us to investigate whether enhanced expression of retrotransposons by the combination of PRC2 loss and DNMTi therapy mediates the profound cytotoxicity upon DNMTi therapy. Retrotransposons, including ERVs, LINE, SINE, can form dsRNAs that are recognizable by various cytosolic dsRNA sensor proteins including RIG-I-like receptors (e.g., RIG-I, MDA5), Oligoadenylate synthases (OASes) (e.g., OAS1/2/3), or protein kinase R (PKR), which can lead to anti-proliferative/survival outcomes^31^ (**Fig. 6E**). To determine which dsRNA sensor mediates lethality of DNMTi in PRC2-loss cancer context, we engineered knockout of either MAVS (a key signaling adaptor downstream of RIG-I and MDA5)^31^, RNaseL (a key effector activated by OAS1/2/3)^31^, and PKR^31^ in PRC2-isogenic cells (**Fig. 6E-F**). Only knockout of PKR was able to partially rescue DNMTi-induced cytotoxicity in PRC2-loss context (**Fig. 6G**).

**Figure 6.**
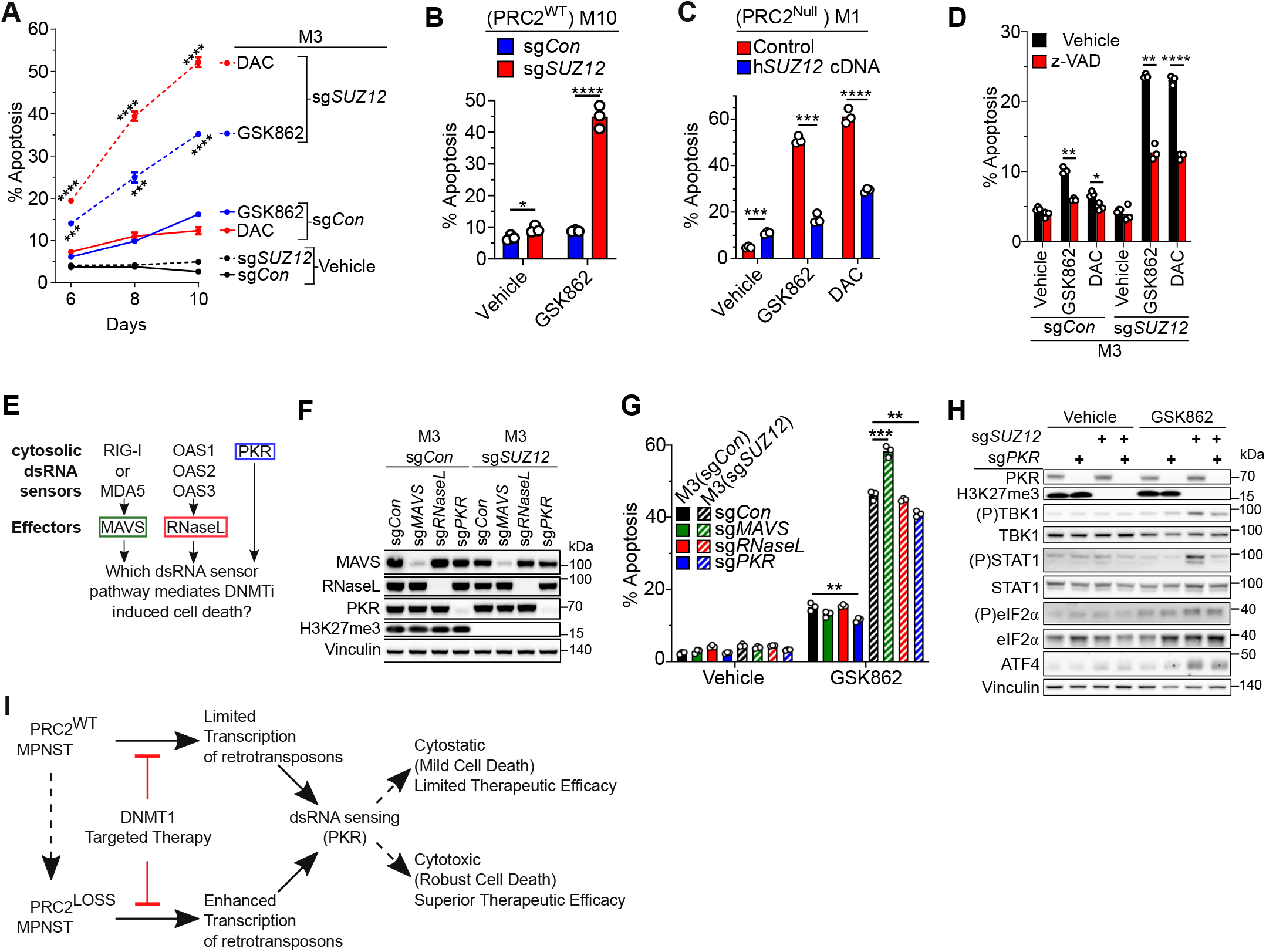
PRC2 inactivation enhances DNMTi-induced programmed cell death via PKR-mediated viral mimicry sensing. **A**, FACS analysis for apoptosis (APC-AnnexinV+) in *SUZ12*-isogenic M3 cells under various treatment conditions. **B**, FACS analysis for apoptosis (APC-AnnexinV+) in DNMTi therapy treated *SUZ12*-wt M10 cells ±sg*SUZ12*. **C**, FACS analysis for apoptosis (APC-AnnexinV+) in DNMTi therapy treated *SUZ12* null M1 cells ±*SUZ12* re-expression. **D**, FACS analysis of DNMTi-induced apoptosis (APC-AnnexinV+) in *SUZ12*-isogenic M3 cells with pan-Caspase inhibitor (z-VAD-FMK) rescue. **E**. A schematic of several main dsRNA sensors and their direct effectors. **F**, Western blot validation of CRISPR/Cas9 mediated knockout of MAVS, RNaseL, and PKR. **G**. Effect of MAVS, RNaseL, or PKR knockout on 3 μM GSK862-mediated apoptosis by APC-AnnexinV FACS analysis. **H**, Western blot of DNMTi-mediated induction of PKR signaling, and attenuation by PKR knockout in *SUZ12*-isogenic M3 cells. **I**, Proposed model of mechanisms underlying enhanced therapeutic efficacy of DNMTi therapy in cancer with PRC2 inactivation. All data presented as the mean ± SEM; Student’s two-tailed T-test (\*\**P*<0.01; \*\*\**P*<0.001; \*\*\*\**P*<0.0001).

Binding of dsRNA to PKR has been reported to cause subsequent downstream phosphorylation of eIF2α, and increased ATF4 protein level^38,39^. Consistently, PKR signaling manifested by downstream signaling effect was further enhanced by DNMTi treatment in PRC2 loss context (e.g., phosphorylation of TBK1, STAT1, and eIF2α, increased ATF4 protein levels); this is partially rescued by PKR knockout, thus corroborating the augmented PKR pathway activation and effects with DNMTi treatment in PRC2 loss context (**Fig. 6H**). Collectively, these findings suggest that loss of PRC2 creates a specific therapeutic vulnerability to DNMTi through augmented expression of retrotransposons and subsequent PKR activation (**Fig. 6I**).

## DISCUSSION

Despite the clinical success of targeting PRC2 in PRC2-dependent cancers^40,41^, therapeutically targeting tumor suppressor loss in PRC2-loss cancers remains challenging. Through systematic RNAi screening in this study, we identified DNMT1 mediated DNA methylation as an epigenetic synthetic lethal interaction with PRC2 inactivating mutations in MPNST. Loss of either one of the PRC2 core components (SUZ12, EED, or EZH2) is sufficient for this synthetic lethal interaction, which suggests a critical role for PRC2 histone methyltransferase activity for the phenotype. In support of this, our integrative genomic analyses revealed dual regulation of a subset of H3K27me3 gene targets by PRC2 and DNMT pathways. Specifically, upon PRC2 loss in MPNST, DNMT1 mediated DNA methylation is associated with restricted expression of a subset of H3K27me3 gene targets while transcription of non-DNMT regulated H3K27me3 gene targets are robustly activated by PRC2 loss alone. These findings are consistent with human embryonic stem cells in which loss of TET enzymes leads to acquisition of promoter DNA methylation at H3K27me3 target genes, which subsequently maintains their silencing after H3K27me3 removal upon cellular differentiation^42^. PRC2 can function as an oncogenic driver or tumor suppressor depending on the cancer cell context. Given that PRC2 is a bona-fide tumor suppressor in MPNST pathogenesis^9-11^, our data may suggest that DNA methylation restricts transcription of subset of H3K27me3 gene targets that are anti-fitness in tumorigenesis and point to potential therapeutic opportunities.

Decitabine and 5-Azacytidine have also been shown to induce innate immune signaling and modest anti-proliferative effects via activation of the RIG-I/MDA5-dependent dsRNA sensing, and subsequent viral mimicry in various solid tumor cell lines without particular molecular features^28,29^. Although HDACi and DNMTi have been shown to have anti-proliferative effects in various cancer cell lines *in vitro*^37,43-45^, these agents have limited clinical activity as a single agent, likely due to the lack of cytotoxic effects in the non-selective contexts of solid tumors^37,45,46^. Our study through non-biased RNAi-based screen identified DNMT1 as a synthetic lethal interaction candidate with PRC2-loss cancer context. Using the pan-DNMTi (at low doses that minimizes off-target effect) and a novel DNMT1-selective small molecule inhibitor^23^, we demonstrated that the DNMTi can induce robust caspase-dependent programmed cell death *in vitro* and potent anti-tumor effect *in vivo* in the PRC2-loss cancer context, which suggests a potential therapeutic window. We further demonstrated that DNMTi cooperatively induces augmented expression of various retrotransposon elements, including ERV/LTRs, LINEs, SINEs, which in turn activates the proinflammatory and innate immune signaling responses probably through multiple dsRNA sensing mechanisms (e.g., RIG-I/MDA5, PKR, OASs’, etc). However, for the DNMTi-mediated programmed cell death effects in the PRC2-loss context, PKR, but not RIG-I/MDA5/MAVS or OAS/RNaseL, mediated dsRNA sensing and activation of downstream signaling is required for robust anti-tumor effects, in contrast to previous studies^28,29,47^. It is worth noting that different dsRNA sensors can have differential RNA substrate binding specificity based on dsRNA features, including length, secondary structures, local mismatches, noncanonical pairs etc^31,48^. Understanding how those features dictate which dsRNA sensing pathway mediates an anti-tumoral outcome in response to DNMT inhibitors may be warranted in future investigations.

Our study provides the scientific rationale for clinically evaluating DNMT inhibitors in patients with PRC2-loss MPNST that currently have an unmet clinical need. To date, the majority of preclinical therapeutic candidates for MPNST have been derived by insights from PRC2-wt genetically engineered murine MPNST models and/or limited human MPNST preclinical *in vivo* models^49-59^. Our study leveraged multiple authenticated *in vitro* and *in vivo* human patient-derived models of PRC2-wt and PRC2-loss MPNST as well as murine models to identify and validate DNMT1 pathway inhibition as a potential therapeutic strategy for PRC2-loss MPNST. Several pan-DNMT inhibitors, e.g., Decitabine, 5-azacitidine, oral combination of decitabine and cedazuridine, have been FDA approved for treatment of myelodysplastic syndromes (MDS)^22,60,61^. Based on our data, we have just started an investigator-initiated trial of ASTX727 (oral combination of decitabine and cedazuridine) in patients with PRC2-loss MPNST (ClinicalTrials.gov, NCT04872543). Ideally, clinical evaluation of DNMT1 specific inhibitors that minimize the off-target effect should be considered in cancers with PRC2 loss context. An orally bioavailable DNMT1 selective inhibitor has recently been shown in murine models to effectively reduce DNA methylation and induce tumor regression while being well tolerated without adverse hematological toxicity^23^, which may circumvent the known side effects of myelosuppression of decitabine^22^. Historically, pan-DNMTs have limited activity in solid tumors due to lack of cytotoxic effects and predictive therapeutic biomarkers^62^. Our study identified the PRC2-loss cancer context as a selective therapeutic vulnerability to DNMTi. PRC2-loss can be readily identified by IHC staining of H3K27me3 and FDA-approved molecular biomarker testing assays, e.g., Foundation One (Foundation Medicine, Inc) or MSK-IMPACT^63^, which may be useful for biomarker-driven clinical investigation of pan-DNMT and DNMT1 targeted therapeutics in solid tumors.

## METHODS

### Cell lines

NF1 patient derived human MPNST cell lines M14 (ST88-14), M3, and M10 are kind gifts developed by Jonathan Fletcher, and William Gerald respectively. *SUZ12* isogenic M10 and M3 MPNST cells were generated by CRISPR/Cas9 targeting of *SUZ12* followed by isolation and validation of single cell clones. At least five single cell clones of sg*Con* or sg*SUZ12* were respectively pooled to create cell lines for experiments. *Eed* isogenic mouse MPNST cells were generated as follows: *Eed*^*f/f*^ mice^64,65^ were bred with PLP-CreERT::*Nf1*^f/f^;*Cdkn2a*^f/f^ mice. *Nf1*^−/–^;*Cdkn2a*^−/–^;*Eed* ^f/–^ cells were cultured from lumbar nerves in tamoxifen treated PLP-CreERT::*Nf1*^f/f^;*Cdkn2a*^f/f^;*Eed*^f/f^ mice. *Eed* null cells were generated by infection with adenovirus-Cre *in vitro. Eed* isogenic cells were infected with sgRNA lentiviruses for additional knockout of *Cdkn2b* followed by subcutaneous implantation into CB-17 SCID mice (Taconic), and upon malignant tumor formation, *Eed* isogenic murine MPNST primary cell lines (*Nf1* ^−/–^;*Cdkn2a* ^−/–^;*Cdkn2b* ^−/–^) were derived from tumors. Histopathology was verified by C.R.A.. HEK-293T were purchased from (ATCC). All cell lines were cultured in DMEM:F-12 (1:1) high glucose media (10% FBS + 15 mM HEPES + 2 mM L-glutamine + 1% penicillin/streptomycin) prepared by the Media Preparation Facility (MSKCC). Exception: MPNST cell lines were cultured in Advanced DMEM/F12 media (Gibco) supplemented with 15% FBS, 2mM L-glutamine, and 1% penicillin/streptomycin for shRNA library screening. Culture conditions (37°C in a 5% CO_2_ incubator). All cell lines tested negative for mycoplasma.

### Virus packaging and infection

Lentiviral transfer plasmid plus packaging plasmids psPAX2 (gift from Didier Trono, Addgene Plasmid #12260, RRID:Addgene_12260) and pCMV-VSV-G (gift from Bob Weinberg, Addgene Plasmid #8454, RRID:Addgene_8454) were co-transfected intro HEK 293T cells via X-tremeGene 9 reagent (Millipore Sigma #6365787001). Two days later, viral media is filtered through a 0.45 μm sterile syringe filter, and stored in-80°C. For infection, thawed virus was mixed with culture media +8 μg/mL Polybrene (Millipore Sigma #107689) to infect cells overnight, followed by removal of virus media. Virus infected cells were selected with appropriate antibiotic after two days post infection.

### shRNA library screening

A doxycycline inducible pooled human epigenome focused lentiviral shRNA plasmid library in the LT3GEPIR mir-E backbone (**Fig. 1B, Supplementary Table 2**)^66^ targeting 565 genes (5 shRNAs per gene) was obtained from the Gene Editing and Screening Core Facility (MSKCC). HEK 293T cells were co-transfected with packaging plasmids to produce lentivirus for shRNA library. Target cell lines were infected with library lentiviruses at ∼10% transduction efficiency followed by selection with 2 μg/mL puromycin for 4 days, and maintenance/expansion of cells at 3000x shRNA representation under 1 μg/mL puromycin for an additional 7 days. For screening, cells were maintained at minimum of 1000x shRNA representation. shRNAs were induced with 1 μg/mL doxycycline, and GFP(+) shRNA expressing cells were isolated by fluorescence activated cell sorting at passage doubling 1 and 15 followed by genomic DNA isolation with Gentra Puregene cell kit (Qiagen). Sequencing libraries were prepared by subjecting 18 μg (1.2 μg x 15 reactions) of genomic DNA (∼1000x shRNA representation) to PCR based barcoding and amplification with AmpliTaq Gold PCR kit (ThermoFisher Scientific). See Supplementary Table 3 for list of barcode adaptor primers. Parallel PCR reactions were pooled, and purified by a column-based PCR purification kit (Qiagen) followed by 30 minute incubation at 37°C with Exonuclease I (New England BioLabs) to digest single stranded

DNA, subsequent separation by agarose gel electrophoresis, and gel extraction via Gel Purification Kit (Qiagen). Purified libraries (PCR product) were subjected to single-end, 50 base-pair high-throughput sequencing on an Illumina HiSeq2500 platform. All resulting FASTQ files passed quality control by FastQC^67^. Sequencing reads were trimmed with Cutadapt^68^ to keep only 22-mer shRNA guide sequences that were subsequently mapped to shRNA library guide sequence reference file using bowtie2^69^. shRNA read counts were quantified by edgeR^70^, Ranking of each set of 5 shRNAs per gene in the library was ranked for depletion with HiTSelect^71^. See **Supplementary Table 1** for resulting shRNA ranking data.

### Drug and chemicals

See Supplementary Table 4 for Drugs, Chemicals, and Regents used in this study. Cells were treated with low dose Decitabine for 4 consecutive days in all *in vitro* experiments.

### Dose response and cellular growth/viability assays

ATP CellTiter-Glo 2.0 cell viability assays (Promega) were performed as previously described^55,72^ to generate dose response and growth curves in Figure 2 and Extended Data Figure 2. All data plotted with GraphPad (Prism).

### Animal studies

Animal experiments were carried out in accordance with protocols approved by the MSKCC Institutional Animal Care and Use Committee (IACUC), and were in compliance with relevant ethical regulations regarding animal research. For DAC studies on human MPNST xenograft bearing CB-17 SCID mice (Taconic): all cells were implanted into mice 1:1 with Matrigel (BD Biosciences). M3 cells (stably infected with MSCV-Luciferase-PGK-Hygro retrovirus) were orthotopically transplanted into the sciatic nerve pocket of mice, and resulting tumors were verified (histopathology by C.R.A.) and re-grafted into new mice for DAC studies when tumors reached steady state growth. M14 cells (stably infected with MSCV-Luciferase-PGK-Neo-IRES-GFP retrovirus) were orthotopically transplanted into the sciatic nerve pocket of mice. The resulting tumors were verified (histopathology by C.R.A.), and tumorigenic cell lines were established. These cells were then re-injected into mice for DAC studies when tumors reached steady state growth. MPNST PDX-1 tumors in CB-17 SCID mice were enzymatically dissociated followed by subcutaneous injection into flanks of new CB-17 SCID mice. When tumors reached 50-100 mm^3^ on average, mice were separated into Vehicle (PBS) and DAC treatment groups based on similar representation of tumor sizes and mouse weights. Mice were treated with Vehicle (PBS) or 5 mg/kg DAC (as previously described^73^) by intraperitoneal injection once daily for 3 consecutive days followed by no further treatment for at least 11 days. As previously described^23^, 45 mg/kg GSK3685032 or vehicle was formulated and administered by intraperitoneal injection twice daily to mice. M3 and M14 xenograft tumor growth was measured by luciferase bioluminescence imaging as previously described^72^. Tumor volume was calculated with the following formula [(4/3)π x (length/2) x (width/2) x (depth/2)] using caliper-based measurements of each tumor. For inducible *DNMT1* knockdown xenografts studies, M14 tumorigenic cells (Luciferase+) were stably infected with doxycycline inducible shRNAs (without doxycycline addition), and orthotopically transplanted into the sciatic nerve pocket of CB-17 SCID mice (Taconic). For *in vivo* knockdown, mice were fed doxycycline water (200 mg/L + 0.5% sucrose). Xenograft tumor growth was measured by luciferase bioluminescence imaging as previously described^72^. All data plotted with GraphPad (Prism).

### FACS analysis

All flow cytometry was performed using LSRFortessa Flow Cytometer (BD Biosciences). GFP linked shRNA competition assays were performed as described previously^74^. Apoptosis was assayed using APC-Annexin V (Tonbo biosciences) or Annexin V-FITC Kit (Miltenyi Biotec). All data were processed and analyzed with FCS Express 7 Research Edition (De Novo Software, Inc). All data plotted with GraphPad (Prism).

### DNA cloning

Control (sh*Ren*.713), *DNMT1, DNMT3A*, and *DNMT3B* shRNAs were cloned into Tet-ON all-in-one plasmids LT3GEPIR (gift from Johannes Zuber, Addgene Plasmid #111177, RRID:Addgene_111177) and LT3REPIR (same as LT3GEPIR except for GFP being substituted with dsRed, MSKCC Gene Editing & Screening Core Facility) as previously described^66^. sgRNAs: Control and *SUZ12* sgRNA targeting the VEFS functional domain were cloned into pL-CRISPR.EFS.Blast (plasmid was generated by replacing tagRFP in Addgene Plasmid #57819 with blasticidin S deaminase antibiotic selection marker) as previously described^75^. Human *SUZ12* cDNA was cloned into pLV-EF1α-IRES-Puro (gift from Tobias Meyer, Addgene Plasmid #85132, RRID:Addgene_85132). *MAVS* sgRNA was cloned into LRG2.1_Puro (gift from Christopher Vakoc, Addgene Plasmid #125594, RRID:Addgene_125594)^76^. Control, RNaseL and PKR sgRNAs were cloned into lentiCRISPR v2 (gift from Feng Zhang, Addgene Plasmid # 52961, RRID:Addgene_52961). See Supplementary Table 5 for shRNA and sgRNA target sequences.

### RNA extraction, reverse transcription and quantitative PCR (qRT-PCR)

Total RNA was isolated from cell lines via TRIzol reagent (ThermoFisher Scientific). For tissues, samples were disrupted with TissueRuptor II (Qiagen) in TRIzol reagent followed by RNA isolation. Total RNA (1 μg) was used for cDNA synthesis with High-Capacity cDNA Reverse Transcription kit (ThermoFisher Scientific). cDNA and primers were mixed with PowerUp SYBR green 2X master mix (ThermoFisher Scientific), followed by mRNA quantification with QuantStudio 6 Flex Real-Time PCR System (Applied Biosystems). *RPL27* was used as the housekeeping gene for *in vitro* cells while human specific *GAPDH* was used as the housekeeping gene for human xenograft tissue samples. Relative fold change gene expression was calculated by the delta-delta Ct method. All data plotted with GraphPad (Prism). See Supplementary Table 6 for qPCR primers used.

### Western blot

Cells were lysed with RIPA buffer (ThermoFisher Scientific) supplemented with protease and phosphatase inhibitors (Millipore Sigma) followed by brief sonication (Diagenode Bioruptor), and quantified by BCA assay (ThermoFisher Scientific). Whole cell lysates were mixed with NuPAGE LDS sample buffer (ThermoFisher Scientific) supplemented with 2-mercaptoehtanol, incubated at 95°C for 10 minutes, and loaded onto NuPAGE 4-12% Bis-Tris gels (ThermoFisher Scientific) for SDS-PAGE followed by protein transfer onto 0.2 μm nitrocellulose membranes (Cytiva Life Sciences) by wet electroblotting at 4°C. Membranes were blocked for 1 hour at room temperature with StartingBlock TBS buffer (ThermoFisher Scientific), and incubated with primary antibody at 4°C overnight with gentle rotation. Next day, membranes were washed thrice with 1x TBS-T, and incubated with HRP-conjugated secondary antibody for 2 hours at room temperature with gentle rotation. Membranes were washed thrice with 1x TBS-T followed by application of SuperSignal West Pico PLUS chemiluminescent substrate (ThermoFisher Scientific) or Immobilon Western chemiluminescent HRP substrate (Millipore Sigma) to membranes, and chemiluminescence scanning with ImageQuant LAS4000 (GE Healthcare). See Supplementary Table 7 for antibodies used. Resulting data was processed and cropped with GNU Image Manipulation Program (GIMP) open-source software.

### Targeted Next Generation Sequencing Assay

Memorial Sloan Kettering-Integrated Mutation Profiling of Actional Cancer Targets (MSK-IMPACT)^63^ was used to evaluate mutations for 341 cancer genes in human MPNST cell lines.

### RNA sequencing and data analysis

*SUZ12* isogenic M3 MPNST cells were treated in triplicate with or without DNMT inhibitors (50 nM daily Decitabine or 4 μM single dose of GSK862) for 4 days followed by total RNA isolation. Library preparation and sequencing was conducted by the Integrated Genomics Operation (IGO) core facility at MSKCC. After RiboGreen quantification and quality control by Agilent BioAnalyzer, 500ng of total RNA with RIN values of 8.3-10 underwent polyA selection and TruSeq library preparation according to instructions provided by Illumina (TruSeq Stranded mRNA LT Kit, catalog # RS-122-2102), with 8 cycles of PCR. Samples were barcoded and run on a HiSeq 4000 in a PE50 run, using the HiSeq 3000/4000 SBS Kit (Illumina). An average of 41 million paired reads was generated per sample. Ribosomal reads represented 2% of the total reads generated and the percent of mRNA bases averaged 76%.s

RNA sequencing reads were 3’ trimmed for base quality 15 and adapter sequences using version 0.4.5 of TrimGalore (https://www.bioinformatics.babraham.ac.uk/projects/trim_galore), and then aligned to human genome assembly hg19 with STAR v2.6 using default parameters. Data quality and transcript coverage were assessed using the Picard tool CollectRNASeqMetrics (http://broadinstitute.github.io/picard/).

TPM data was generated via TPMCalculator using default parameters. TPM data for C1, C4, and C6 gene were plotted on violin plots using GraphPad (Prism). Read count tables were generated with HTSeq v0.9.1.

Normalization and expression dynamics were evaluated with DESeq2 using library size factor normalization and outliers were assessed by sample grouping in principal component analysis. These library size factors were also used with deepTools v3.1 to create normalized bigwigs using bamCoverage with --scaleFactor.

All differentially expressed genes (FDR < 0.1 and log_2_ fold change > 1.5), from pairwise analyses were then merged to create a union of all contrasts, and *k*-means clustering was performed from *k*=4 to the point at which cluster groups became redundant. Pathway enrichment (gene sets and gene ontology) was performed on each cluster using Homer v4.5 (http://homer.ucsd.edu) and comparing with the cumulative hypergeometric distribution.

For differential expression analysis of retrotransposon elements, we extracted LTRs, SINEs and LINEs from the repetitive elements (REs) annotated by the RepeatMasker and used featureCounts (v1.5.0-p1)^77^ to count RNA-seq fragments mapped to them. Note that RNA-seq reads mapped to multiple regions were kept by STAR. REs with a minimal 10 reads in the RNA-seq samples were retained for differential expression analysis using DESeq2 with default parameters (v1.33.4). Significantly differentially expressed REs (padj < 0.1 and fold change > 1.5) in at least one of our six comparisons were subject to *k*-means clustering via pheatmap (v.1.0.12), separately for the three types of REs.

### Bisulfite sequencing and data analysis

*SUZ12* isogenic M3 MPNST cells were treated with or without DNMT inhibitors (50 nM daily Decitabine or 4 μM single dose of GSK862) for 4 days followed by genomic DNA isolation with Gentra Puregene cell kit (Qiagen). Bisulfite sequencing was conducted by the IGO core facility at MSKCC. Briefly, after PicoGreen quantification and quality control by Agilent BioAnalyzer, 500 ng of genomic DNA were sheared using a LE220-plus Focused-ultrasonicator (Covaris catalog # 500569).

Sequencing libraries were prepared using the KAPA Hyper Prep Kit (Kapa Biosystems KK8504) without PCR amplification. Post-ligation cleanup proceeded according to Illumina’s instructions with 110 µL Sample Purification Mix from the TruSeq Methyl Capture EPIC LT Library Prep Kit (Illumina catalog # FC-151-1002). After purification, 4 samples were pooled equivolume and methylome regions were captured using EPIC oligos. Capture pools were bisulfite converted and amplified with 11 cycles of PCR. Pools were sequenced on a HiSeq 2500 in Rapid mode in a 100bp/100bp paired end run, using HiSeq Rapid SBS Kit v2 (Illumina). The average number of read pairs per sample was 61 million. Adapter trimming, alignment to hg19, and CpG methylation calling was done as previously described^78^. *methylKit*^79^ R package was used to filter all samples for CpGs with 10-400x coverage followed by tiling the genome into 1000bp windows (regions) with 1000bp step-size, and identification DMRs (> +/- 25% change). DMRs were annotated via *annotatr* ^*80*^ R package. For intergenic DMRs, the nearest gene was identified by the *closest* function in BEDTools suite^81^. Genomation^82^ R package was used to annotate CpGs and tiles(regions) with their location (CpG island, shores, promoters), and then plotted as violin plots via ggplot2^83^ package in R. Principal component analysis of methylation data at all tiles (regions) was generated in Partek Genomics Suite (v 7.0). For DNA methylation profile plots, 4 individual replicates (PBS rep1 and 2, DMSO rep1 and 2) were averaged for each cell line followed by visualization of the mean DNA methylation profile over regions of interest via plotProfile command from deepTools ^84^. CGI locations were obtained from UCSC genome table browser, and used for extracting promoter CGIs for subsequent visualization of CpG DNA methylation as density plots or quantification by plotHeatmap and multiBigwigSummary commands respectively from deepTools^84^.

### Chromatin immunoprecipitation (ChIP) sequencing and data analysis

Recipe for ChIP related buffers can be found in Supplementary Table 8. *SUZ12* isogenic M3 MPNST cells were treated with or without DNMT inhibitors (50 nM daily Decitabine or 4 μM single dose of GSK862) for 5 days. Cells were harvested followed by cross-linking with 1% formaldehyde for 10 minutes at room temperature, quenched with glycine (125 mM final). Cells were washed twice with sterile cold PBS, followed by lysis with LB1 buffer (+protease and phosphatase inhibitors for 10 minutes at 4°C while rotating, pelleted and resuspended in LB2 buffer (+protease and phosphatase inhibitors) for 10 minutes at 4°C while rotating, and pelleted/resuspended in SLB buffer (+protease and phosphatase inhibitors) for 10 minutes at 4°C while rotating. Lysates were diluted 10-fold with ChIP dilution buffer (+protease and phosphatase inhibitors) followed by chromatin shearing with a Covaris E220 Focused Ultrasonicator (20% Duty Cycle + 150W PIP + 200 cycles/burst). Sheared chromatin was centrifuged for 10 minutes at 4°C, and supernatant was collected followed by addition of antibody + magnetic protein A/G beads for overnight rotation at 4°C. See Supplementary Table 7 for antibodies used. Next day, beads were washed on a magnetic stand thrice with cold High Salt Wash buffer, and once with Low Salt Wash buffer. Immunoprecipitated chromatin DNA was isolated from beads with Elution buffer at 65°C for 30 minutes. Cross links were reversed overnight at 65°C. ChIP DNA was diluted 2-fold with ChIP TE+NaCl buffer, and incubated 2 hours at 37°C with RNaseA followed by incubation at 55°C for 2 hours with Proteinase K. ChIP DNA was purified by PCR purification kit (Qiagen). Samples were submitted to the IGO core facility at MSKCC for library construction, and sequencing. Briefly, pooled multiplexed libraries were sequenced on a HiSeq 2500 machine on rapid mode at 50bp paired-end reads per sample.

Adaptor sequences were removed from resulting data with TrimGalore followed by alignment to human genome assembly hg19 using bowtie2 (v2.3.5) with default parameers^69^. Duplicates reads were removed. Coverage bigwig files were created via bamCoverage command from deepTools ^84^ to normalize to total reads and human genome size, and subsequent ChIP-seq profiles were visualized by either Integrative Genomics Viewer^85^ software or by using the plotHeatmap command from deepTools^84^. Genomic regions enriched with H3K27me3 reads were identified using a sliding window method, using input as controls as described previously^86,87^. Genes with peaks in their promoter regions (+/2 kb from TSS) were designated as H3K27em3+ genes.

### Statistical analysis

All experiments (excluding high throughput sequencing) were repeated at least twice independently. All error bars were shown as the mean ± SEM. A two tailed student’s T-test was used to evaluate statistical significance (\**P* ≤ 0.05, \*\**P* ≤ 0.01, \*\*\**P* ≤ 0.001, \*\*\*\**P* ≤ 0.0001).

## DATA AVAILABILITY

RNA-seq, bisulfite-seq, and ChIP-seq data have been deposited at GEO under accession codes GSE179585, GSE179582, and GSE179586. Any additional information required to reanalyze the data reported in this paper is available from the lead contact upon request.

## AUTHORS’ CONTRIBUTIONS

Project planning and experimental design (A.J.P., Y.C. and P.C.). Animal experiments (A.J.P, S.W., M.R.P., E.F., D.L., N.W., C.J.L., E.G.). Cell culture (A.J.P., G.B., Y.M.). DNA cloning (A.J.P.). ChIP-seq (A.J.P.). Dose response cell viability assays (A.J.P., M.M-R., G.B.). FACS analysis (A.J.P.). shRNA screening (A.J.P.). qRT-PCR (A.J.P.). Western blots (A.J.P., Y.M.). Technical support: (S.W., E.W.P.W., J.Y., C.J.L., and E.G.). M.B.P and M.T.M for early access/guidance on usage of GSK862 (GSK3484862) and GSK3685032. J.M.S. for guidance on *in vitro* and *in vivo* decitabine usage. Histopathology of tumors (C.R.A.). Analysis of genomics data (A.J.P., J.L.M, R.M., R.P.K., J.L.G., D.Z.). Manuscript writing (A.J.P., Y.C. and P.C.). All authors reviewed and/or edited the manuscript.

## ACKNOWLEDGEMENTS

We thank Drs. William Gerald and Leon (Xiaoliang) Xu (MSKCC) for patient derived M3 cell line. Melissa Pappalardi and Michael McCabe (GlaxoSmithKline plc.) for providing GSK862 and GSK3685032, and advice on their usage. We thank Dr. Scott Armstrong for generously providing *Eed*^*f/f*^ mice^64,65^. We thank Makhzuna Khudoynazarova for help with generating MPNST mouse models and human MPNST cell line derived xenograft model. We acknowledge the use of the Integrated Genomics Operation (IGO) Core, funded by the NCI Cancer Center Support Grant (CCSG, P30 CA08748), Cycle for Survival, and the Marie-Josée and Henry R. Kravis Center for Molecular Oncology. IGO core conducted library preparation and next generation sequencing for ChIP-seq, bisulfite-seq, and RNA-seq samples. Processing and analysis of bisulfite-seq and RNA-seq data by the Center for Epigenetics Research (MSKCC). This work was supported in part by grants from the National Institute of Health (NIH) and National Cancer Institute (NCI) grants (R01 CA228216, DP2 CA174499), Department of Defense (DOD) grant (W81XWH-15-1-0124), Francis Collins Scholar NTAP, Cycle for Survival and Linn Family Discovery Fund to P.C.; the NIH/NCI grant (P50 CA217694) to P.C., C.R.A.; the NIH/NCI grants (5R01CA208100-04, 5U54CA224079-03, 5P50CA092629-20) to Y.C.; Geoffrey Beene Cancer Research Fund to P.C., M.L.; Department of Defense Horizon Award (CA181474) to M.A.M.; Translational Oncology Research in Oncology Training Program T32 grant (5T32CA160001-09) to A.J.P.; NIH grant P30 CA 008748 to Memorial Sloan Kettering Cancer Center (Core Grant).

## FIGURE LEGENDS

**Supplementary Figure S1(Related to Figure 1).**
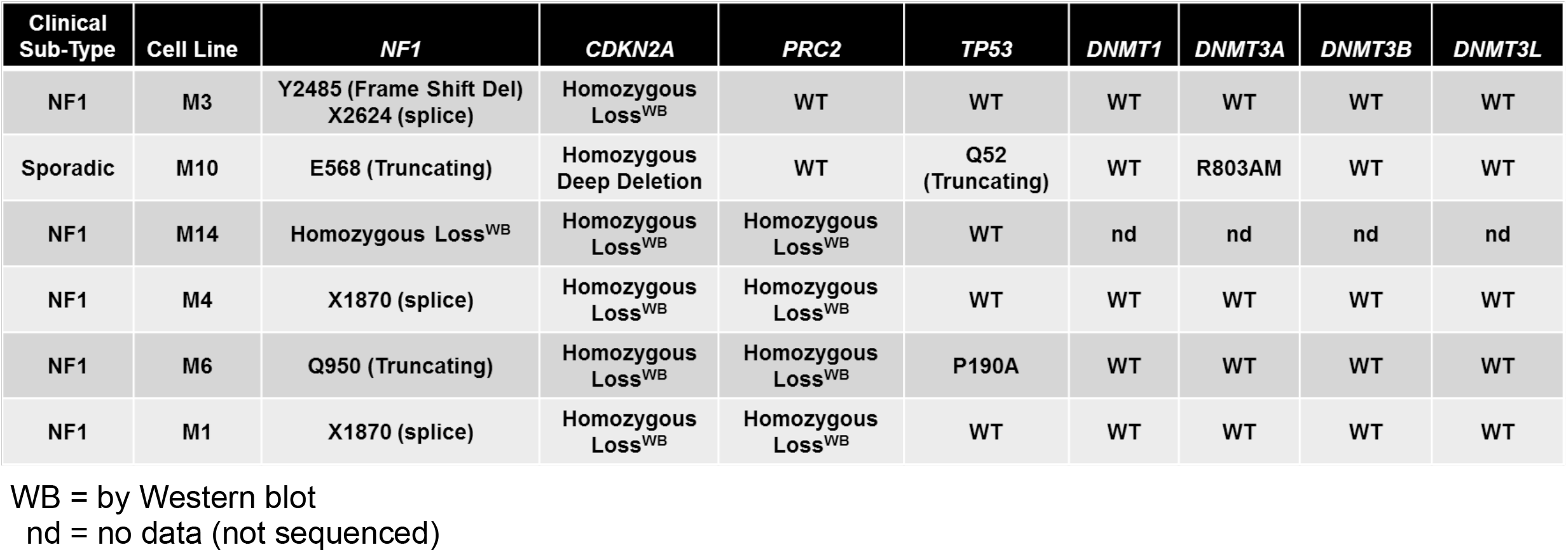
Table of pathologically validated human MPNST cell line information. Genetic alterations are based on targeted next generation sequencing assay (MSK-IMPACT) unless specified otherwise.

**Supplementary Figure S2(Related to Figure 1).**
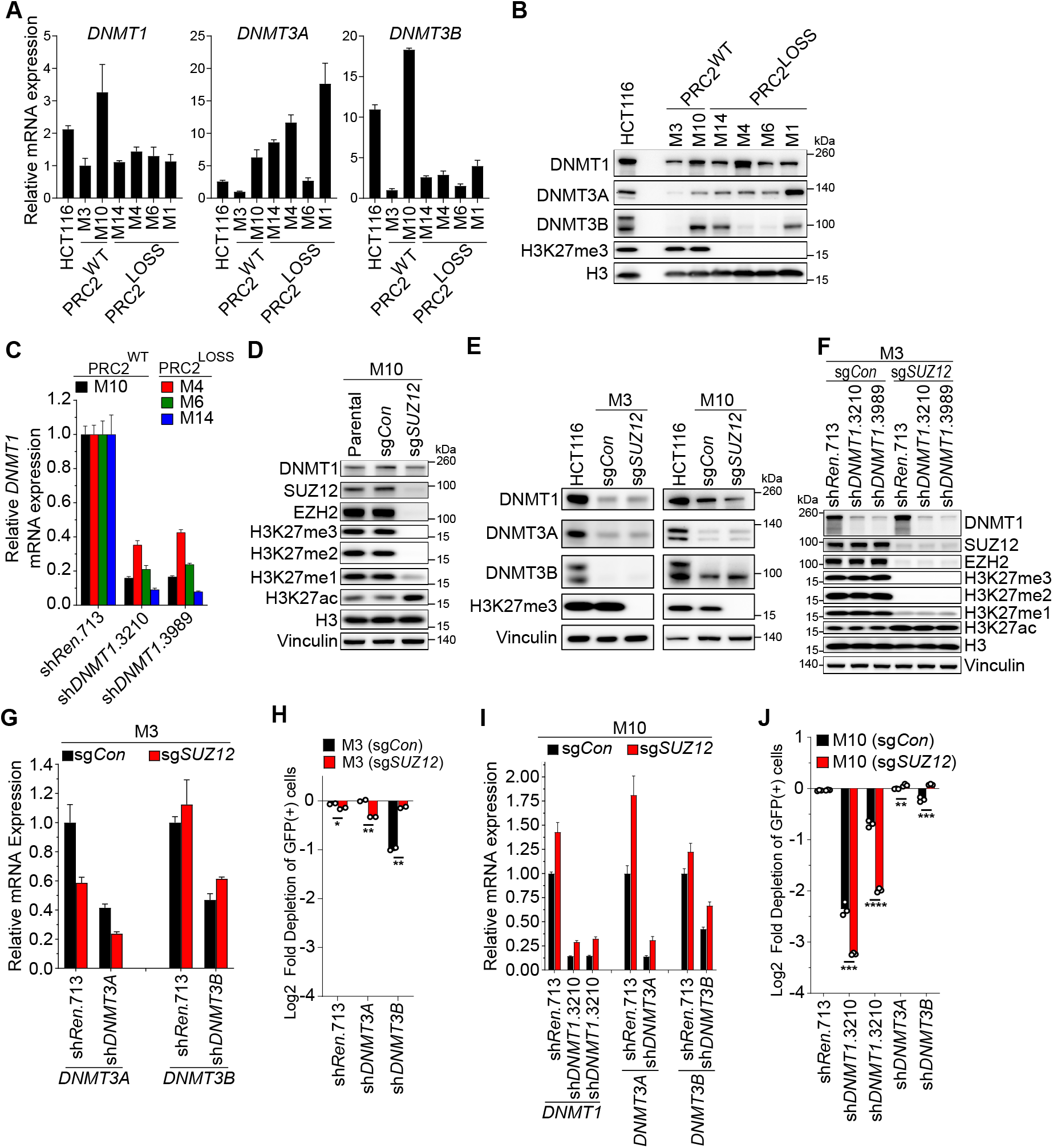
**A-B**, RNA expression by qRT-PCR (**A**) and protein levels by immunoblot (**B**) of DNMTs in various patient-derived PRC2-wt and PRC2-loss MPNST cell lines. **C**, qRT-PCR validation of *DNMT1* mRNA knockdown at 4 days of doxycycline-inducible expression of *DNMT1* shRNAs in non-isogenic human MPNST cell lines. **D**, Western blot validation of *SUZ12*-isogenic M10 cells. **e**, Western blot of DNMT protein expression in a *SUZ12*-isogenic human MPNST cells. **F**, Western blot validation of sgRNA-mediated knockout of *SUZ12* leading to loss of PRC2 function, and doxycycline-inducible knockdown of *DNMT1*. **G**, qRT-PCR validation of doxycycline-inducible knockdown of *DNMT3A* or *DNMT3B* mRNA in *SUZ12*-isogenic M3 MPNST cells. **H**, Effect of *DNMT3A* or *DNMT3B* knockdown in cells via shRNA-linked GFP by FACS-based competition assay in indicated cell lines. **I**, qRT-PCR validation of doxycycline-inducible knockdown of *DNMT1, DNMT3A*, or *DNMT3B* in *SUZ12-*isogenic M10 cells. **J**, Validation of *DNMT1* as a synthetic lethal candidate with PRC2-inactivation compared to *DNMT3A/B* via shRNA-linked GFP by FACS-based competition assay in *SUZ12-*isogenic M10 cells. All error bars: mean ± SEM; Student’s two-tailed T-test (\*\**P*<0.01; \*\*\**P*<0.001; \*\*\*\**P*<0.0001).

**Supplementary Figure S3(Related to Figure 2).**
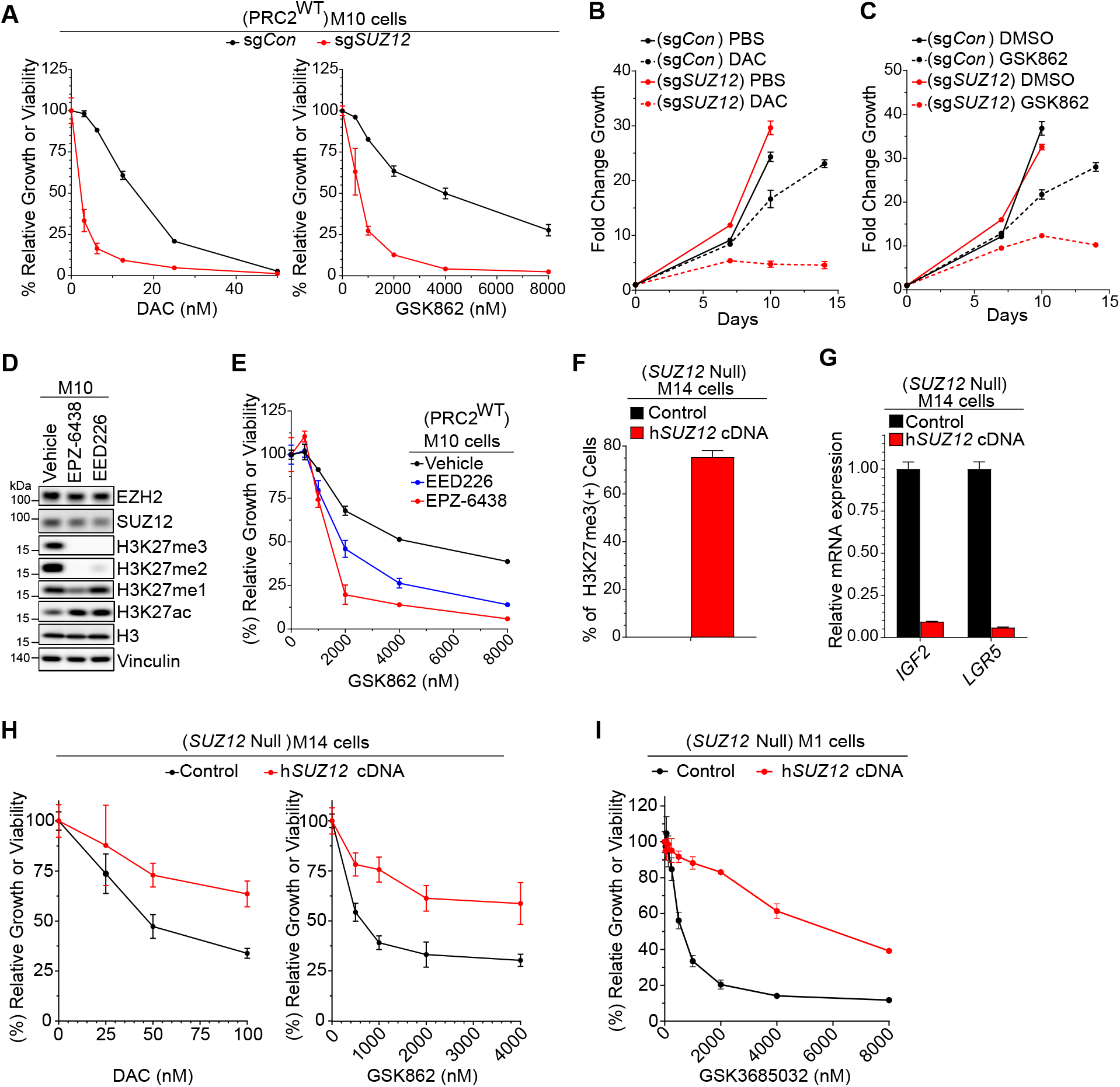
**A**, Dose-response curves of a pan-DNMT inhibitor, decitabine (DAC) or a DNMT1-selective inhibitor (GSK862) in *SUZ12-*isogenic M10: PRC2-wt= sg*Con*; PRC2-loss = sg*SUZ12*. **B-C**, Growth curves of *SUZ12-*isogenic M3 cells treated with 50 nM DAC or 4 μM GSK862. **D**, Western blot analysis of PRC2-wt M10 cells treated with 3 μM EPZ-6438 (EZH2 inhibitor) or 5 μM EED226 (EED inhibitor). **E**, GSK862 dose-response curves for PRC2-wt M10 cells treated with EZH2 or EED inhibitors or vehicle controls. **F**, Percentage of H3K27me3 positive cells assessed by immunofluorescence in parental *SUZ12*-null M14 cells with *SUZ12* restoration. **G**, Relative mRNA expression of PRC2 target genes in *SUZ12* loss MPNST cells with or without *SUZ12* restoration. **H**. DAC and GSK862 dose-response curves in *SUZ12*-null human MPNST cells with or without PRC2 restoration. **I**. GSK3685032 dose-response curves in *SUZ12*-null human MPNST cells with or without PRC2 restoration. All error bars: mean ± SEM.

**Supplementary Figure S4(Related to Figure 3).**
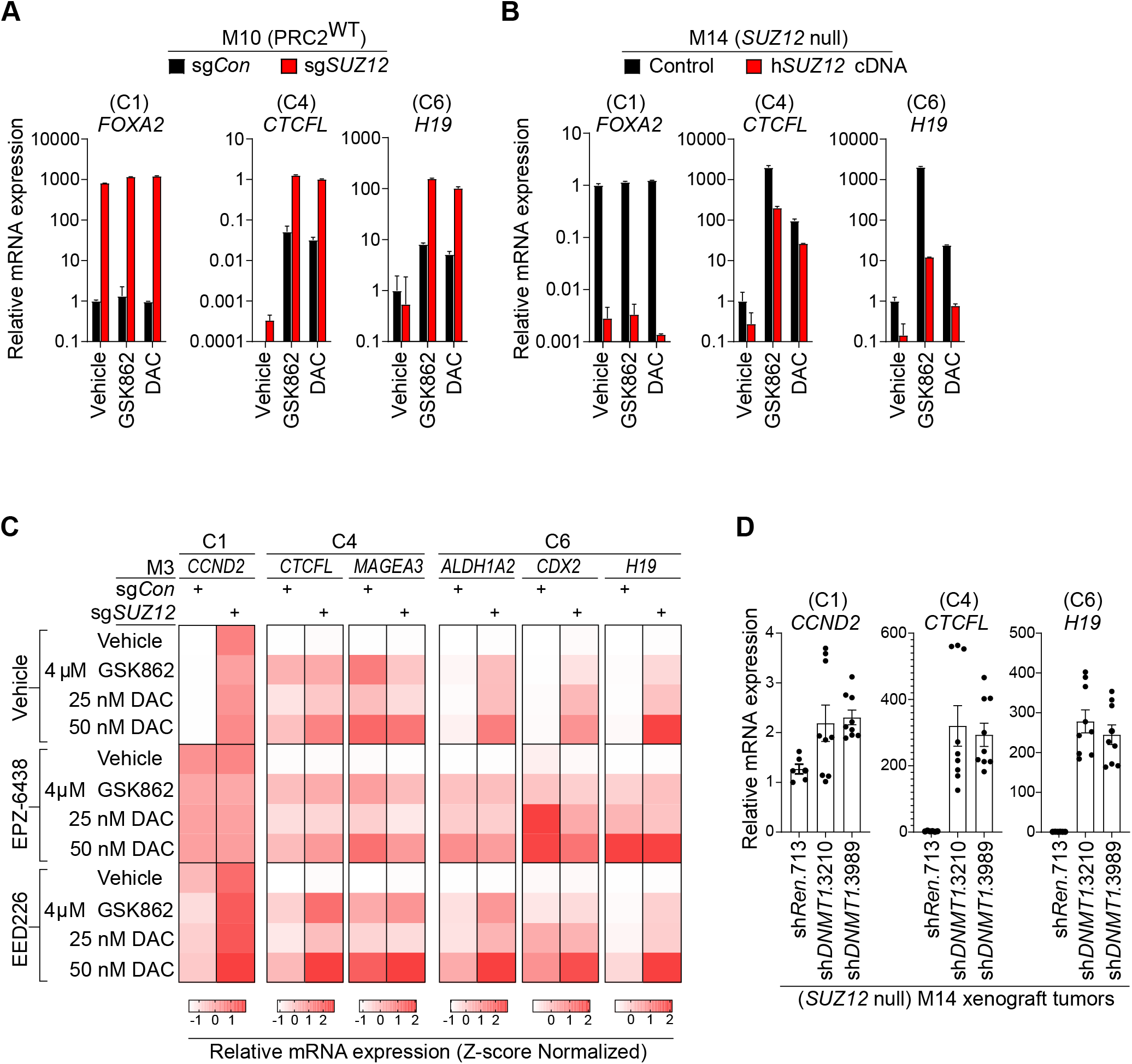
**A-B**, qRT-PCR analysis of representative genes in C1, C4, and C6 in *SUZ12*-isogenic M10 and M14 cells under various treatment conditions. **C**, Heatmap of Z-score normalized mRNA expression of C1, C4, C6 genes by qRT-PCR in PRC2-isogenic M3 cells under various treatment conditions, PRC2 inhibitors (5 μM EPZ-6438, 3 μM EED226) and/or DNMTi (GSK862, DAC). **D**, Relative RNA expression by qRT-PCR for C1, C4, and C6 genes in PRC2-loss M14 xenograft tumors in mice with doxycycline inducible knockdown of *DNMT1*. All error bars: mean ± SEM.

**Supplementary Figure S5(Related to Figure 4).**
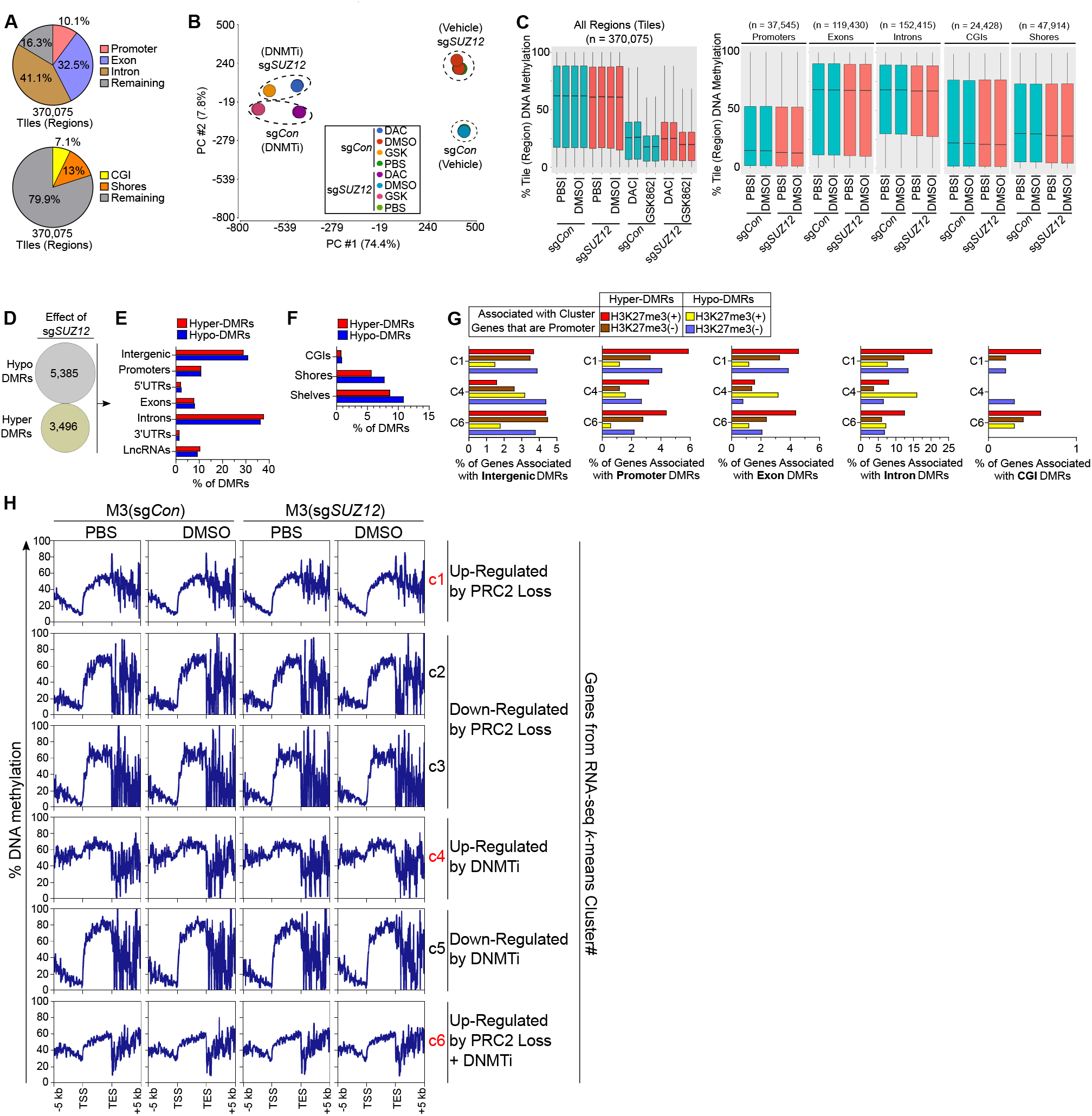
**A**, Genic and CpG feature distribution for genomic tiles (regions) shared between all bisulfite-sequencing samples. **B**, Principal component analysis of tile (region) CpG DNA methylation data for *SUZ12*-isogenic M3 cells under various treatments. **C**, Box plots of DNA methylation levels for all regions, promoters, exons, introns, CGIs, CpG shores in PRC2-isogenic cells under various treatment conditions. **D**, Number of Differentially Methylated Regions (DMRs) caused by PRC2 loss (via sg*SUZ12*). DMR: methylation difference > 25%, *q*< 0.01). **E-F**, Percentage of DMRs associated with genic or CpG features. **G**, Percentage of sg*SUZ12*-mediated DMRs in C1, C4, and C6 genes and separated by H3K27me3 enrichment status (±). **H**, Average DNA methylation profiles spanning the TSS to the TES for C1-6 genes in *SUZ12*-isogenic M3 cells.

**Supplementary Figure S6(Related to Figure 6).**
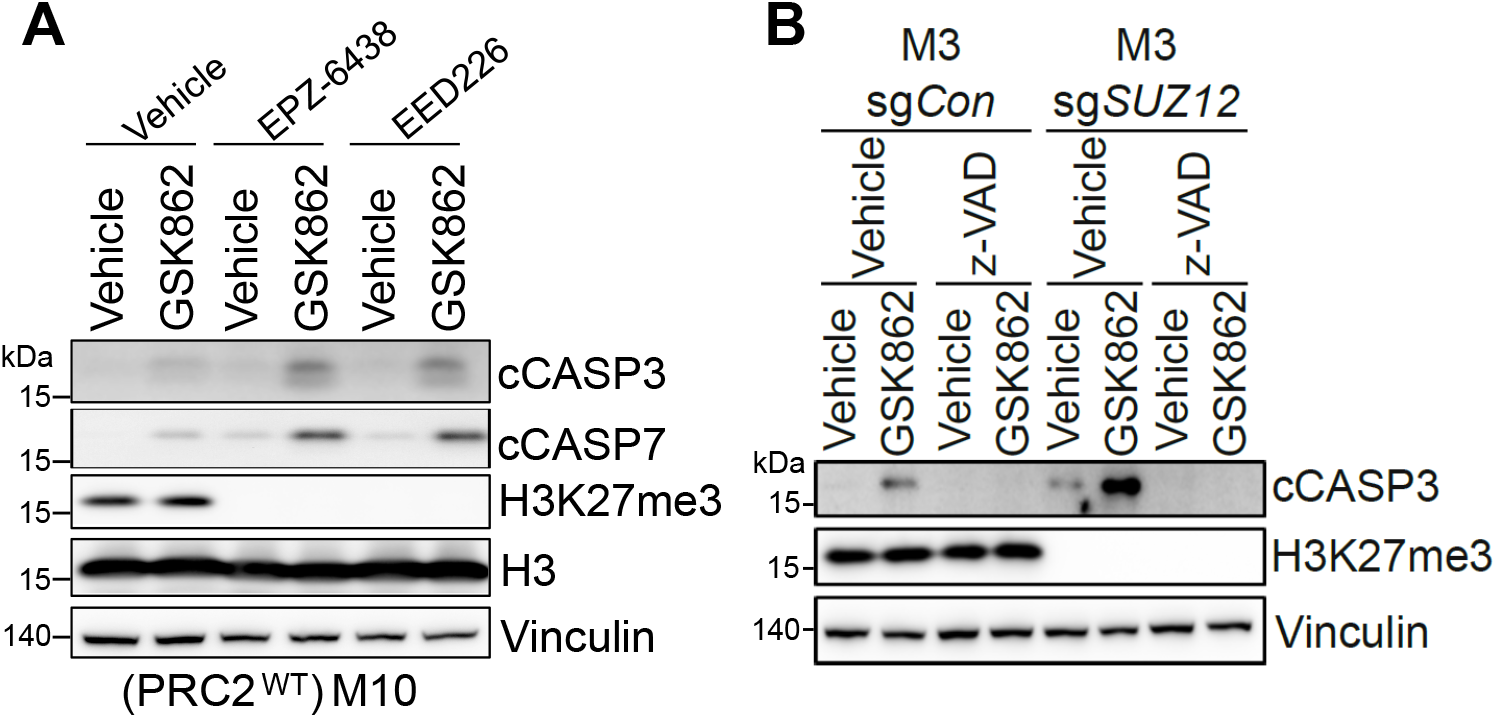
**A**, Western blot of Caspase cleavage in PRC2-wt M10 cells pre-treated with 5 μM EZH2 inhibitor (EPZ-6438) or 3 μM EED inhibitor (EED226) for 10 days followed by ±4 μM GSK862 treatment. **B**, Western blot of Caspase cleavage in PRC2-isogenic M3 cells treated ±4 μM GSK862 ±10 μM z-VAD for 6 days.

